# Start codon-associated ribosomal frameshifting mediates nutrient stress adaptation

**DOI:** 10.1101/2023.02.15.528768

**Authors:** Yuanhui Mao, Longfei Jia, Leiming Dong, Xin Erica Shu, Shu-Bing Qian

## Abstract

A translating ribosome is typically thought to follow the reading frame defined by the selected start codon. Using super-resolution ribosome profiling, here we report pervasive out-of-frame translation immediately from the start codon. The start codon-associated ribosome frameshifting (SCARF) stems from the slippage of ribosomes during the transition from initiation to elongation. Using a massively paralleled reporter assay, we uncovered sequence elements acting as SCARF enhancers or repressors, implying that start codon recognition is coupled with reading frame fidelity. This finding explains thousands of mass spectrometry spectra unannotated from human proteome. Mechanistically, we find that the eukaryotic initiation factor 5B (eIF5B) maintains the reading frame fidelity by stabilizing initiating ribosomes. Intriguingly, amino acid starvation induces SCARF by proteasomal degradation of eIF5B. The stress-induced SCARF protects cells from starvation by enabling amino acid recycling and selective mRNA translation. Our findings illustrate a beneficial effect of translational “noise” in nutrient stress adaptation.

## INTRODUCTION

Eukaryotic translation initiation involves more than a dozen initiation factors orchestrating ribosome loading, scanning, and start codon selection ^1,2^. Although the main start codon almost exclusively establishes the primary open reading frame (ORF), frameshifting occurs during elongation either spontaneously or in a sequence-programmed manner ^3,4^. Another mechanism contributing to out-of-frame translation is alternative initiation using start codons in different reading frames ^5^. Since non-AUG codons could serve as potential initiation sites ^6^, it is not always apparent whether downstream out-of-frame translation is due to elongation-associated frameshifting or alternative initiation. The key mechanistic difference lies in the fidelity of start codon recognition, as start codon skipping leads to “leaky scanning” and downstream initiation. Once a start codon is recognized by the scanning ribosome, the engagement of the initiator tRNA is followed by 60S joining, a process facilitated by the evolutionarily conserved GTPase eIF5B ^7^. While our knowledge about start codon selection is steadily increasing, the transition of the assembled 80S from an initiation complex to an elongation complex remains incompletely understood ^8^. Neither the timescale nor the quality control of the 80S assembly at the start codon is known prior to elongation commitment.

Ribosome profiling (Ribo-seq) is a powerful technique that provides a snapshot of global translation by sequencing ribosome-protected mRNA fragments (RPFs) ^9,10^. Notably, Ribo-seq revealed substantial amount of out-of-frame footprints ^11,12^, but their origins remain largely ambiguous. We do not know exactly, beyond a few examples, whether the off-track translation is a result of alternative initiation, frameshifting during elongation, or simply experimental noise. Mass spectrometry (MS) is the main methodology that enables direct identification of translational products. Surprisingly, on average, 75% of spectra analyzed in an MS experiment cannot be identified ^13^. Since search engines were built upon theoretical spectra derived from user-defined protein sequences, it is possible that the current proteome database is far from complete. Even when all three reading frames are considered, the identity of millions of spectra remains elusive. Without knowing the scope of translational diversity, how translational “noises” contribute to the proteome landscape remains a fundamental knowledge gap.

Here, we re-designed the Ribo-seq methodology by introducing <u>e</u>a<u>sy</u> <u>R</u>NA-<u>a</u>denylation <u>seq</u>uencing (Ezra-seq), which not only simplifies library construction but also improves the quality of Ribo-seq. With superior resolution, Ezra-seq revealed pervasive out-of-frame translation after the main start codon. We interpretated these translational “noises” as a result of start codon-associated frameshifting (SCARF). Importantly, we uncovered a regulatory role for eIF5B in controlling SCARF, suggesting a quality control mechanism in maintaining start codon-associated reading frame fidelity. Intriguingly, nutrient stress induces SCARF via eIF5B degradation, implying physiological significance of translational “noise”.

## RESULTS

### Ezra-seq offers high resolution ribosome profiling

Typical Ribo-seq results are characterized by the 3-nt periodicity of ribosome-protected mRNA fragments (RPFs). The percentage of in-frame reads (IFR), or phasing, is often used to gauge the quality of Ribo-seq data sets. However, different methods result in varied 5’ end read accuracy. For instance, the standard Ribo-seq approach relies on circularization after reverse transcription that is known to introduce untemplated nucleotide addition ^14^. RNA ligation method suffers from sequence-dependent biases ^15^, resulting in altered footprint quantification. Additionally, the varied read length often requires offset adjustment to infer the correct ribosome P-site position ^14^. Despite continuous optimization of Ribo-seq methodology, a substantial number of reads remain out-of-frame (Extended data Fig. 1a). We re-designed the Ribo-seq methodology by introducing <u>e</u>a<u>sy</u> <u>R</u>NA-<u>a</u>denylation <u>seq</u>uencing (Ezra-seq) (Extended data Fig. 1b). Without 3’ end linker ligation and 5’ end circularization, Ezra-seq simplifies the library construction and, importantly, improves the 5’ end accuracy of RPFs with > 90% of in-frame reads (Fig. 1a). Furthermore, the ribosome position inferred from the 5’ end of footprints is independent of the read length (Fig. 1a, bottom panel), offering direct determination of P-site positions without *in silico* adjustment. As expected, Ezra-seq revealed prominent peaks around start and stop codons corresponding to initiating and terminating ribosomes, respectively (Extended data Fig. 1c). The single nucleotide resolution of Ezra-seq is highly reproducible in different cell types (Extended data Fig. 1d).

**Fig. 1.**
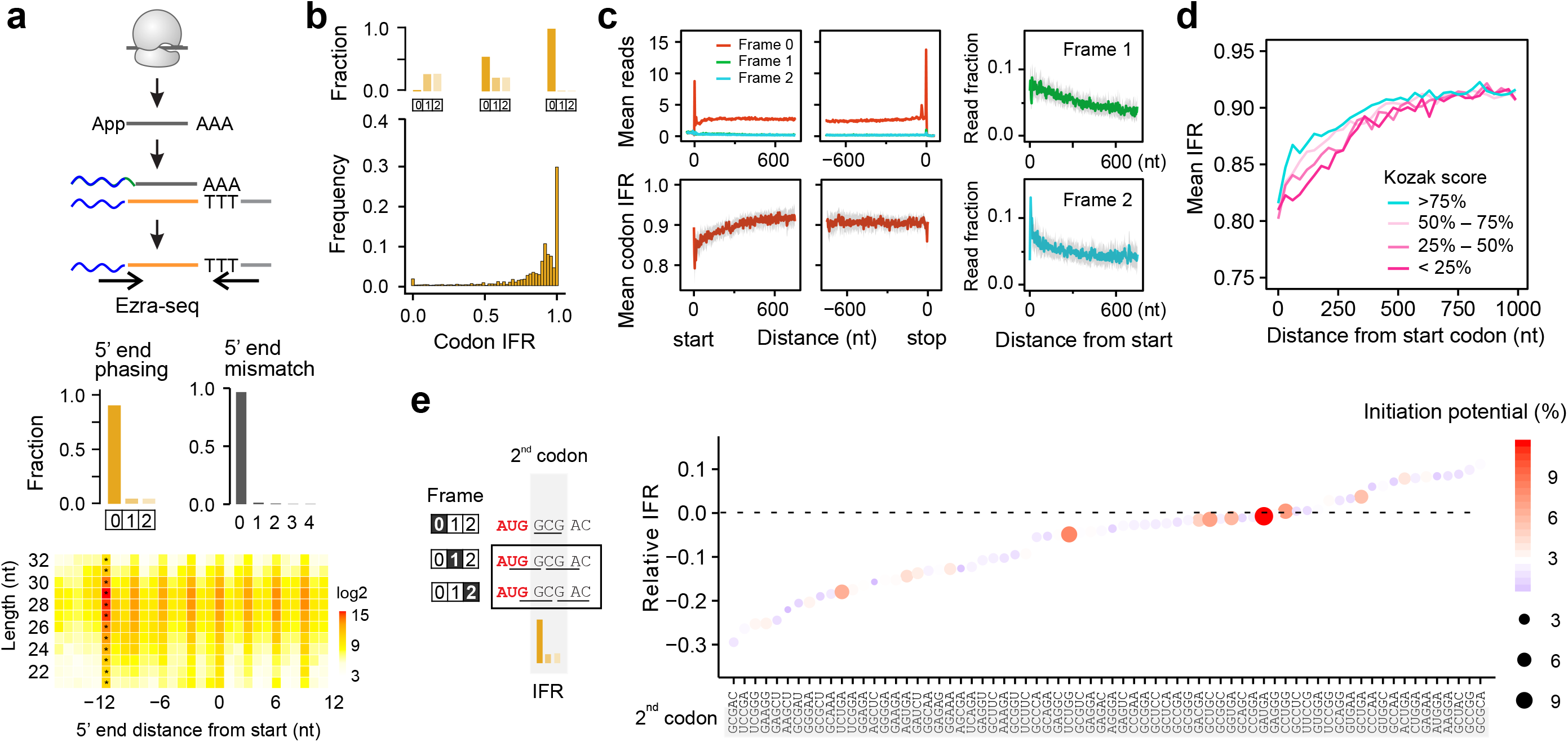
Ezra-seq reveals prevailing out-of-frame footprints in the beginning of CDS. **(a)** The top panel shows schematic of Ezra-seq procedures. The middle panel shows the fraction of reads in different reading frames (5’ end phasing) and the fraction of reads with varied number of 5’ end mismatches. The bottom panel is the heatmap showing the distance of 5’ end to the start codon (*x*-axis) for reads with different length (*y*-axis). The color code represents the log2 read count. **(b)** The top panel shows examples of codons with varied 5’ end phasing corresponding to different codon in-frame ratio (IFR). The bottom panel shows the frequency of codons with different IFR with 0.0 indicating no in-frame reads and 1.0 indicating complete in-frame reads. **(c)** Aggregation plots show the ribosome density (top) and in-frame ratio of ribosome footprints (bottom) across the transcriptome. Right panel shows the fraction of out-of-frame reads across the transcriptome. Grey shadow shows the variation of mean IFR estimated by bootstrap method. Transcripts are aligned to start and stop codons, respectively. **(d)** Comparison of IFR between mRNAs bearing start codons with varied strength of the Kozak sequence context. IFR values are calculated by dividing the in-frame reads by total reads within a non-overlapping sliding window (30 nt). **(e)** Scatter plot shows relative IFR values of the 2^nd^ codons subtracted from the IFR values of the same codons in CDS excluding the first 600 nt. The 2^nd^ codon together with two downstream nucleotides are shown. Color and size of the dots represent the out-of-frame initiation potential around the 2^nd^ codon by averaging initiation potential of 4 out-of-frame codons.

To affirm the superior resolution of Ezra-seq, we conducted a direct comparison using multiple Ribo-seq data sets obtained using different methods (Extended data Fig. 2a – 2h). Ezra-seq clearly offers the highest resolution of footprints as evidenced by increased 5’ end phasing and diminished 5’ end mismatch. Since a start codon is followed by increased in-frame reads downstream (Extended data Fig. 2i), the improved resolution of Ezra-seq allows us to evaluate the initiation potential of all 64 codons. This analysis revealed additional 10 triplets besides AUG capable of initiation as evidenced by >5% increase of IFR for downstream regions. The initiation potential of non-AUG codons is highly conserved across different cell types and species (Extended data Fig. 2j). This result is not only consistent with previous profiling of initiating ribosomes ^16^, but also in line with the measurement of translation initiation in *E. coli* using reporter-based assays ^17^.

### Characterizing out-of-frame footprints

Although out-of-frame reads are substantially reduced by Ezra-seq, the remaining “noisy” footprints are expected to be distributed evenly across individual codons. To our surprise. while ~30% of codons contain perfect in-frame reads, many codons are deprived of in-frame footprints (Fig. 1b). The wide range of IFR across individual codons argues against the possibility of technical bias. Intriguingly, metagene analysis revealed an accumulation of out-of-frame reads in the beginning of the coding region (CDS) (Fig. 1c, right pane). Accordingly, the codon IFR averaged across the transcriptome shows a sharp drop (~20%) following the start codon. This feature is irrespective of cycloheximide treatment and even discernable from regular Ribo-seq results (Extended data Fig. 3a). We surveyed published Ribo-seq data sets and found that the poor codon IFR after the start codon is a common feature across different cell lines (Extended data Fig. 3b). If out-of-frame reads represent footprints of actively translating ribosomes, the density of these reads is expected to drop at the out-of-frame stop codons. This is indeed the case (Extended data Fig. 3c), suggesting that the observed out-of-frame reads are footprints of actively translating ribosomes.

Besides the asymmetric distribution of out-of-frame footprints, we observed large IFR variations across individual transcripts (Extended data Fig. 4a). Transcripts with differential IFR values exhibit similar 5’UTR properties such as folding potential, GC content, and length (Extended data Fig. 4b). Upstream open reading frames (uORFs), when overlapping with the main CDS, would result in out-of-frame footprints after the main start codon. However, a direct comparison between transcripts with or without uORF translation showed comparable IFR patterns in the beginning of CDS (Extended data Fig. 4c). Since most uORFs use non-AUG initiators, it is possible that the overall uORF translation is not robust enough. Indeed, only the overlapping uORFs with the AUG initiator exhibit the lowest IFR (Extended data Fig. 4d). To further exclude the possibility of hidden uORFs, we selected transcripts with an out-of-frame stop codon before the main start codon (e.g., <u>UG**A**</u>**UG**). By excluding overlapping uORFs, those mRNAs still show an evident drop of IFR in the beginning of CDS (Extended data Fig. 4e).

Another possibility is leaky scanning, which often occurs when the start codon is suboptimal ^18^. Indeed, mRNAs bearing weak start codons (without the Kozak sequence context) tend to have lower IFR values in the beginning of CDS (Fig. 1d), although a strong start codon is still associated with a substantial number of out-of-frame footprints. Downstream initiation relies on the presence of downstream AUG (dAUG) triplets, however, most dAUG and cognate sites (dNUG) are > 6 nt away from the main start codon (Extended data Fig. 4f). Given the possibility of alternative initiation at non-AUG codons, we surveyed the initiation potential of entire out-of-frame codons immediately downstream of the annotated start codons (Fig. 1e). This analysis revealed that most 2^nd^ codons with poor IFR values possess negligible out-of-frame initiation potential. For a sequence (<u>CAU</u>GA) with the strongest out-of-frame initiation potential (i.e., AUG), the IFR value at the 2^nd^ codon <u>CAU</u> was barely changed. We conclude that the prevailing out-of-frame footprints after the main start codon is due to mechanisms beyond leaky scanning.

### Codon optimality contributes to out-of-frame footprints

Many organisms, including human, tend to have poorly adapted codons enriched in the beginning of CDS (Fig. 2a) ^19^. It is possible that non-optimal codons promote ribosomal frameshifting (FS) and subsequent out-of-frame translation. We examined the reading frame fidelity of footprints aligned to individual codons. Intriguingly, the A-site codon optimality is positively correlated with IFR (*p* = 0.003) (Fig. 2b, top panel, and Extended data Fig. 5a). Two non-optimal serine codons AGC and AGU exhibit the lowest IFR values. No such correlation exists when the P-site codon identity is considered (Fig. 2b, right panel). The same feature holds true when uORF-containing mRNAs are not considered (Extended data Fig. 5b). The relaxed reading frame fidelity when a non-optimal codon enters the A-site suggests that delayed tRNA delivery induces FS. The A-site codon-induced FS is likely influenced by the P-site codon identity ^3^. Indeed, pair-wise codon analysis revealed that certain triplets at the P-site, such as the tryptophan codon UGG, coordinate with the non-optimal codons at the A-site in lowering IFR (Extended data Fig. 5c).

**Fig. 2.**
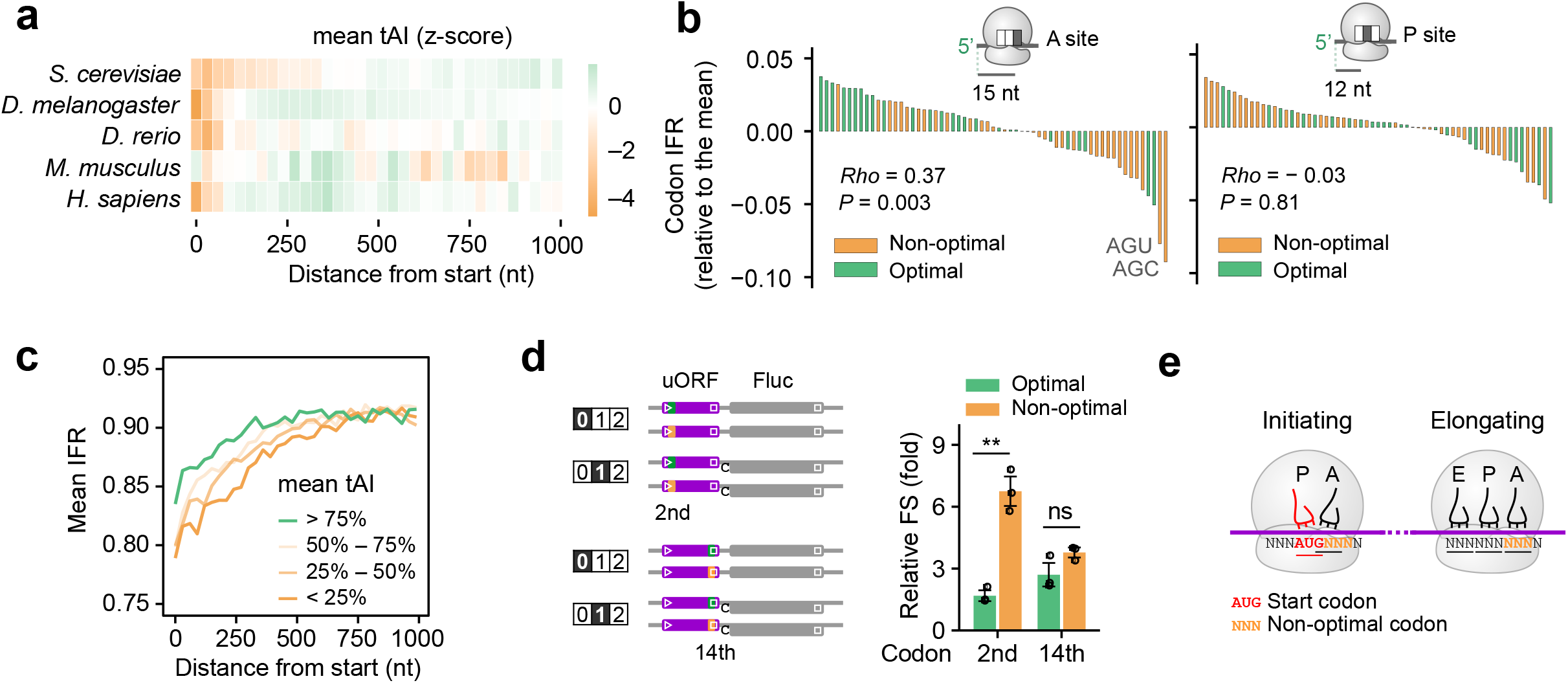
Non-optimal codons induce ribosome frameshifting. **(a)** Heat maps showing mean tAI values along the CDS across different species. tAI values are calculated as a geometric mean of tAI value of the codons in a non-overlapping sliding window (30 nt) along CDS. To compare codon optimality across species, the tAI values were normalized by the z-score normalization. **(b)** Bar plots showing in-frame ratio of footprints aligned to individual codons at ribosome A-site (left panel) or P-site (right panel). Optimal and non-optimal codons are defined based on relative tRNA abundance (tAI) values. **(c)** Comparison of IFR between mRNAs enriched with differential codon optimality at the first 100 codons. IFR values are calculated within a non-overlapping sliding window (30 nt). **(d)** The left panel shows the schematic of uORF reporters with either the 2^nd^ codon or 14^th^ codon replaced by a synonymous optimal (green) or non-optimal (orange). The highlighted numbers in frames indicate the reading frame of Fluc relative to uORF. The right bar graph shows the relative frameshifting rates determined by the downstream Fluc activities (frame 1/frame 0). Error bars, mean ± s.e.m.; two-tailed t-test, *n* = 3, ** *P* < 0.01. The positions of start and stop codons are indicated by tringles and squares, respectively. **(e)** Differential frameshifting susceptibility between initiating and elongating ribosomes in response to non-optimal codons at the A site.

By grouping transcripts with differential codon optimality in the beginning of CDS, we found that the presence of non-optimal codons indeed lowered IFR values in a dose-dependent manner (Fig. 2c). Of note, for many transcripts, the strongest codon bias occurs near the start codon. For instance, the non-optimal alanine codon GCG is mainly found at the 2^nd^ codon position, but barely present in the remaining CDS (Extended data Fig. 5d). To examine the positional effect of codon optimality, we devised a reporter assay by placing the firefly luciferase (Fluc) into different reading frames relative to an uORF (Extended data Fig. 5e). We eliminated out-of-frame stop codons from the uORF and introduced optimal or non-optimal codons without altering the encoded amino acid sequence. To avoid transcription-associated variation, we synthesized mRNA reporters and monitored Fluc levels in transfected HEK293 cells. Presence of the non-optimal UUG at the 2^nd^ codon position resulted in >3 fold higher FS than the optimal CUG (Fig. 2d). Importantly, the same codon swapping at the end of the uORF (14^th^ codon) minimally triggers FS, suggesting that initiating ribosomes are more susceptible to frameshifting than elongating ribosomes (Fig. 2e).

### Start codon-associated ribosome frameshifting

Since the 80S ribosome assembled at the start codon has empty E and A sites ^16^, we propose that the initiating ribosome is susceptible to start codon-associated ribosome frameshifting (SCARF). To test this possibility, we constructed SCARF reporters to monitor the translation of uORF placed in different reading frames (Fig. 3a). The uORF-encoded tracer peptide SIINFEKL is readily presented by the mouse MHC class I molecule H-2K^b^, and the amount of peptide-K^b^ complex can be quantified by a monoclonal antibody 25D1 ^20,21^. When the uORF is placed in different reading frames relative to the first AUG, any 25D1 signals must come from SCARF because neither leaky scanning nor elongation-associated frameshifting would produce the fulllength peptides. Once again, we used mRNA transfection to exclude transcription-associated variation. Compared to the GFP control that showed background levels of 25D1, SCARF reporters with out-of-frame uORFs clearly showed elevated 25D1 signals in transfected HEK293 cells expressing H-2K^b^ (Fig. 3a).

**Fig. 3.**
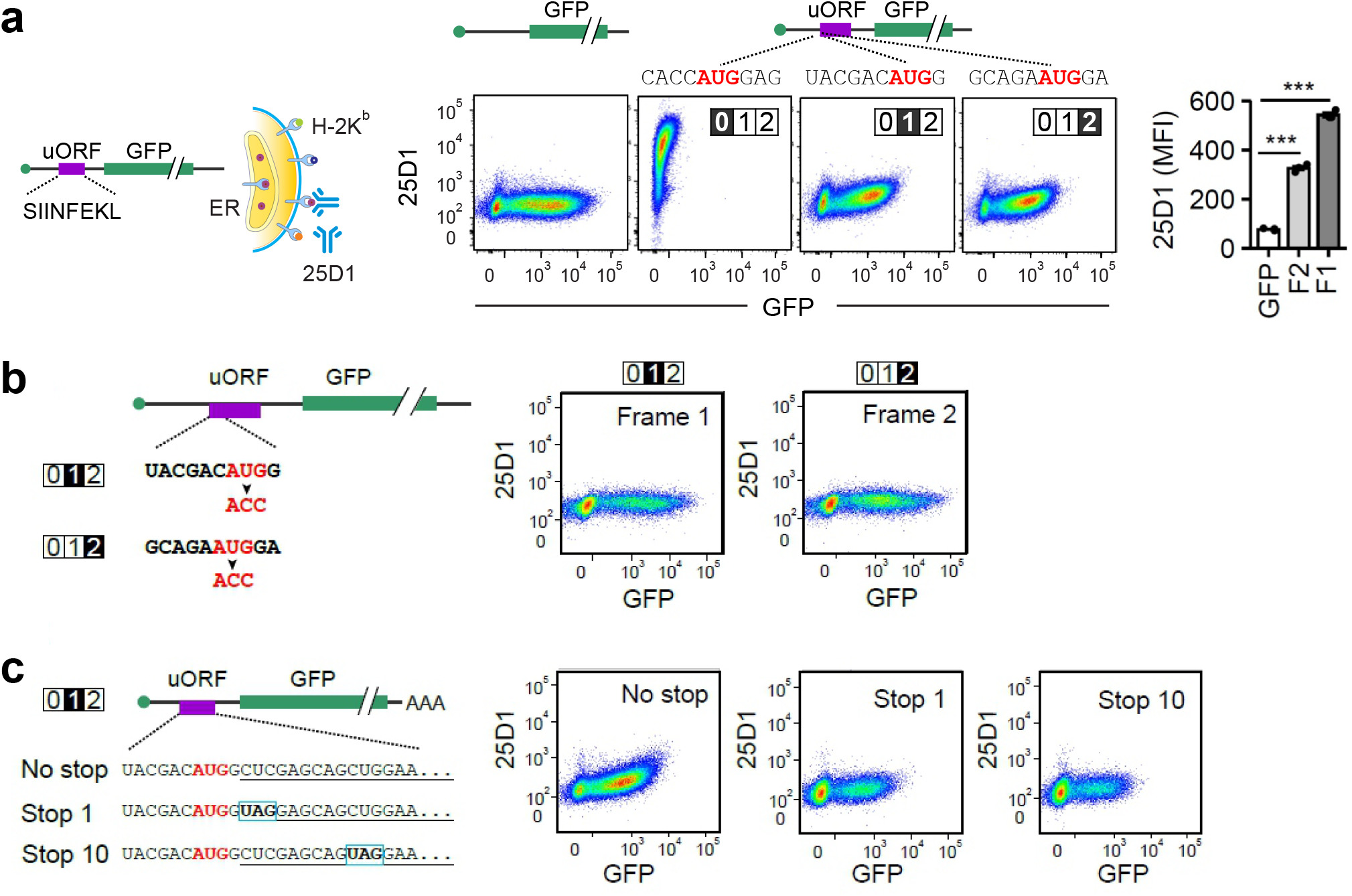
Characterizing start codon-associated ribosome frameshifting. **(a)** The top panel shows the schematic of SCARF reporters measuring start-codon associated frameshifting. The encoded tracer peptide SIINFEKL presented by H-2K^b^ can be quantified by the 25D1 antibody using flow cytometry. The bottom panel shows representative flow cytometry scatter plots of HEK293-K^b^ cells transfected with SCARF reporters as well as GFP control. The highlighted numbers in frames refer to the reading frame of the encoded SIINFEKL relative to the AUG codon. The bar graph shows the 25D1 mean fluorescence intensity (MFI) of SCARF reporters as well as GFP control. Error bars, mean ± s.e.m.; twotailed *t*-test, *n* = 3, *** *P* < 0.001. **(b)** The left panel shows the sequence context of the SCARF reporters with start codon AUG mutated to ACC. The right panel shows representative flow cytometry scatter plots of HEK293-K^b^ cells transfected with SCARF reporters shown on the left. **(c)** The left panel shows the sequence context of the SCARF reporters with an internal stop codon in-frame with SIINFEKL. The position of stop codons is indicated by a frame. Of note, the region encoding SIINFEKL (highlighted by underlines) is placed to reading frame 1 relative to the start codon (highlighted in red). The right panel shows representative flow cytometry scatter plots of HEK293-K^b^ cells transfected with SCARF reporters shown on the left.

Although the SCARF reporter enables us to rule out leaky scanning, it is still possible that SIINFEKL peptides are produced via alternative initiation upstream. We devised two additional reporters to exclude this possibility. First, we mutated AUG to ACC to eliminate the main start codon (Fig. 3b), which would not affect upstream hidden start codons, if any. Lack of AUG abolished 25D1 signals, suggesting that SCARF is coupled with the main start codon. Second, we introduced a stop codon in frame 1 between the start codon and the SIINFEKL coding sequence (Fig. 3c). This design also eliminated the 25D1 signals from transfected cells, suggesting an immediate frameshifting event. Therefore, SCARF represents a previously unrecognized phenomenon by enabling out-of-frame translation from the same start codon.

### SCARF regulation by sequence contexts

To explore the SCARF-associated sequence contexts in an unbiased manner, we took advantage of our previous data sets using a massively paralleled uORF reporter system ^21^. By replacing the start codon of uORF with a random 10-nt sequence, this system allows us to identify SCARF enhancers and suppressors from over a million sequence variants (× 4^10^) (Fig. 4a). To enrich mRNA variants with uORF translation, we separated mRNA reporters based on the number of associated ribosomes using sucrose gradient. Given the small size of uORF (42 nt), mRNAs with preferential uORF translation are expected to reside in the monosome. When an out-of-frame start codon is present, the mRNA variants tend to be enriched in the polysome owing to the longer coding region (Fig. 4a). This approach allows us to separate different mRNAs transfected into the same cell. As expected, AUG was prominently recovered from mRNAs enriched in the monosome fraction (Extended data Fig. 6a). Intriguingly, mRNAs with AUG placed at out-of-frames showed comparable M/P ratios (Fig. 4a), suggesting that many initiating ribosomes are not fixed to the reading frame set by the initiator AUG.

**Fig. 4.**
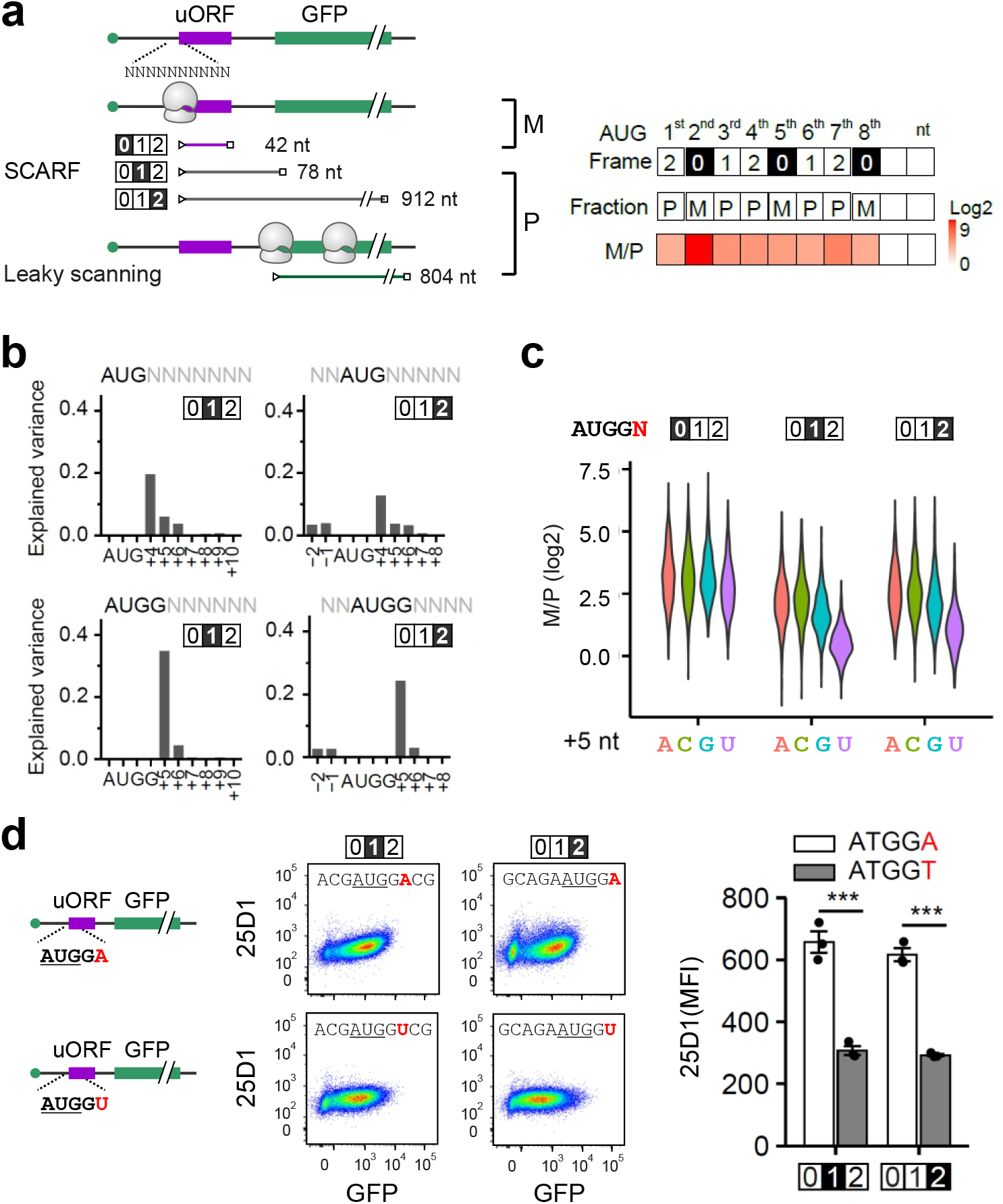
SCARF is sequence context dependent. **(a)** The left panel shows the schematic of a massively parallel SCARF reporter assay, where the start codon of uORF is replaced with a random 10-nt sequence. The uORF translation is monitored by the number of associated ribosomes separated by sucrose gradient. The right panel shows the reading frames relative to uORF when AUG was placed to different positions and the expected ribosome fractions (P, polysome; M, monosome). The heatmap shows the M/P ratio when AUG was placed to different positions. **(b)** Relative contributions of the nucleotide identity in different positions to the uORF translation based on the M/P ratio. The highlighted numbers refer to the reading frame of the encoded SIINFEKL relative to the AUG codon. **(c)** A violin plot shows the influence of the +5^th^ nucleotide identity on the M/P ratios when a Kozak initiation context (RNN<u>AUG</u>G) is placed in different frames. The highlighted numbers refer to reading frame of the encoded SIINFEKL relative to the AUG codon. Twosided Wilcoxon test was performed to test the null hypothesis that the +5 U shows equal M/P ratio as the others. For out-of-frame reporters, all *P* < 2.2 × 10^−16^. **(d)** Representative flow cytometry scatter plots of HEK293-K^b^ cells transfected with SCARF reporters bearing a repressor (thymine at 5^th^ position relative to start codon) or enhancer (adenine at 5^th^ position relative to start codon). Bar plots show the relative 25D1 mean fluorescence intensity (MFI) of SCARF reporters. The highlighted numbers refer to the reading frame of the encoded SIINFEKL relative to the AUG codon. Error bars, mean ± s.e.m.; two-tailed *t*-test, *n* = 3, *** *P* < 0.001.

Notably, the AUG recovery within the 10-nt insert is uneven with the highest frequency at the in-frame 2^nd^ position. Unlike the 2^nd^ position that has the fixed −3 nucleotide, the AUG at the 5^th^ position is surrounded by random sequences. This feature allows us to analyze the sequence context in start codon selection. As expected, the recognition of AUG is primarily dependent on the −3 nucleotide with a minor influence from the +4 nucleotide (Extended data Fig. 6b). The same feature holds true for out-of-frame AUG initiators (Fig. 4b, top panel, and Extended data Fig. 6c – 6d), suggesting that SCARF is also influenced by the Kozak sequence context. Intriguingly, when the AUG is within the optimal sequence context (i.e., RNN<u>AUG</u>G), the +5 nucleotide becomes the most influential in the recovery of out-of-frame AUG codons (Fig. 4b, bottom panel). Analysis of the +5 nucleotide identity revealed U as the least contributor to SCARF (Fig. 4c), indicating that <u>AUG</u>GU suppresses SCARF. Strikingly, the initial IFR analysis of Ezra-seq also uncovered U at the +5 nucleotide position for codons with excellent reading frame fidelity (Fig. 1e). As an independent validation, we constructed SCARF reporters with different +5 nucleotides and confirmed that <u>AUG</u>GU largely prevents SCARF induced by <u>AUG</u>GA (Fig. 4d). The unexpected contribution of the +5 nucleotide identity to SCARF expands the role of the sequence context from start codon recognition to reading frame maintenance.

### Detecting endogenous SCARF products

SCARF is expected to increase the proteome diversity as a result of out-of-frame translation from the start codon. To detect endogenous SCARF products on a global scale, we surveyed the Proteomics IDEntifications (PRIDE) database, the world’s largest data repository of MS-based proteomic data ^22^. We first built a human out-of-frame proteome database by including all *in silico* translation events initiated from frame-shifted start codons. Among newly identified 18,058 out-of-frame peptides (Extended Data Table 1), the majority was surprisingly accumulated following the main start codon (Fig. 5a). Supporting their SCARF origin, transcripts containing out-of-frame peptides exhibit lower IFR values after the start codon (Fig. 5b).

**Fig. 5.**
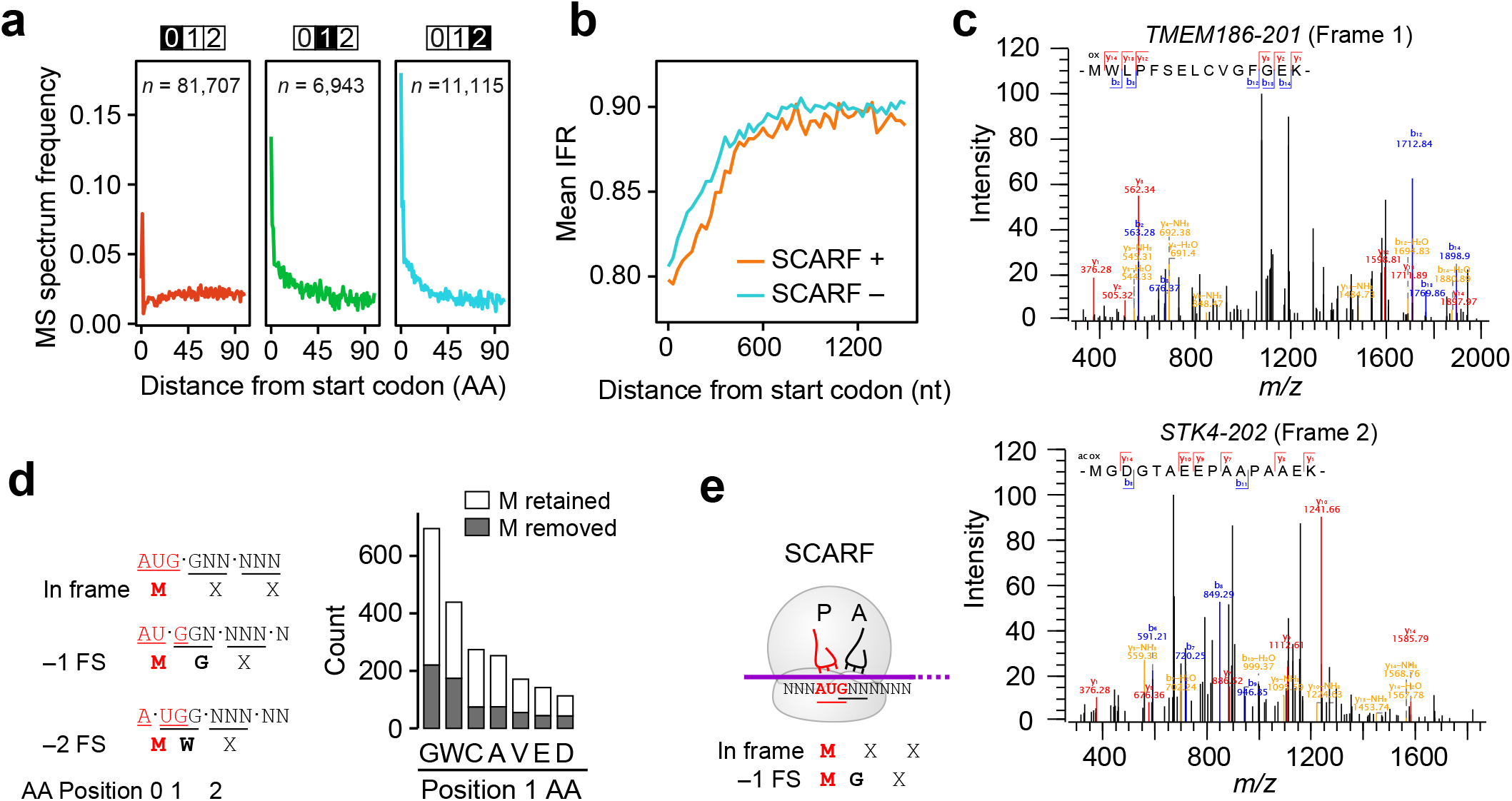
Detecting endogenous SCARF products. **(a)** The frequency of mass spectra mapped to the in-frame and out-of-frame peptides was calculated using the PRIDE database. The out-of-frame spectra were obtained by searching unidentified spectra against the out-of-frame proteome database containing *in silico* translation of SCARF. **(b)** Comparation of IFR between mRNAs with (SCARF +) or without out-of-frame peptides (SCARF -) generated from SCARF. IFR values are calculated within a nonoverlapping sliding window (30 nt). **(c)** Representative examples of mass spectra (An *et al*, 2020) derived from N-terminal out-offrame peptides. **(d)** Count of the N-terminal amino acid after methionine on out-of-frame peptides. M retained, the first methionine is detected at the first N-terminal residue; M-removed, the first methionine is not detected as the first N-terminal residue. **(e)** Schematic of SCARF producing out-of-frame peptides.

One important feature of SCARF is that it commences with the initiator tRNA (Met-tRNAi^Met^) recognizing the main start codon. Therefore, the first amino acid of SCARF peptides should be methionine (M) even though their corresponding ORFs lack the AUG codon. By manually adding an extra M residue at the N-termini (position 0) of out-of-frame peptides, we uncovered 1,403 peptides starting with M (Extended Data Table 1). This number is likely an underestimate because some M residues are removed post-translationally ^23^, leaving a perfect match from position 1 (a total of 685 peptides). Importantly, 62.7% of the extra N-terminal M residues are modified by acetylation, an orthogonal signature of translation initiation ^24^, indicating that they are *bona fide* SCARF products. We next examined whether SCARF peptides could be detected from a single data base. From a multi-plex tandem mass tagging (TMT)-based proteomics study ^25^, we uncovered 167 OFT peptides with a total of 65 derived from the start codon (Fig. 5c and Extended Data Table 2). To substantiate this finding further, we examined a MS dataset encompassing N-terminal peptides enriched by terminal amine isotopic labeling of substrates (TAILS) ^26^. Among 45 OFT peptides, 16 peptides belong to SCARF products (Extended Data Table 3).

Since strong start codons like <u>AUG</u>G contribute to the majority of translational products, we examined the identity of amino acids at position 1 to shed lights on the molecular details of SCARF (Fig. 5d). From the PRIDE database, about a third of the SCARF peptides contain glycine (G) at position 1, an indication of –1 frameshifting because G is encoded by GGN codons; and 21% of peptides contain tryptophan (W), an indication of –2 frameshifting because W is exclusively encoded by UGG (Fig. 5d). These results not only exclude alternative initiation, but strongly suggest that SCARF is due to unstable interaction between the P-site initiator tRNA and the start codon, thereby permitting A-site tRNA mispairing (Fig. 5e).

### SCARF regulation by eIF5B

We next wondered whether SCARF is subjected to regulation. To search for potential SCARF regulators, we focused on initiation factors involved in start codon recognition and 60S joining (Fig. 6a). Upon start codon recognition, eIF1 dissociation enables eIF5 interaction with eIF1A, forming a closed initiation complex to arrest scanning ^27^. Mutants of eIF1 has been shown to reduce the stringency of start codon selection ^28^. However, silencing eIF1 had negligible effect on IFR (Extended Data Fig. 7a), suggesting that SCARF occurs after start codon recognition. Similarly, eIF5 overexpression did not alter the IFR pattern (Extended Data Fig. 7b), albeit compromising the stringency of start codon recognition ^29^. Unlike eIF1 knockdown, eIF1A depletion lowered the IFR immediately after the start codon (Extended Data Fig. 7c). It is worth mentioning that silencing eIF1A led to severe cellular toxicity, presumably due to the pleiotropic functions of eIF1A in nearly all steps of initiation, including 60S joining. The 60S joining is catalyzed by eIF5B ^7^, which also interacts with eIF1A ^30^. When eIF5B was depleted, the IFR values in the beginning of CDS were evidently reduced (Extended Data Fig. 7d). To exclude the possibility that the lowered IFR is an artifactual noise of reduced global translation, we separated mRNAs with differential translation status in cells lacking eIF5B. Remarkably, only mRNAs with high ribosome occupancy experienced reduced IFR (Fig. 6b), indicating that SCARF is coupled with active translation. Supporting the crucial role for eIF5B in start codon-associated reading frame fidelity, SCARF reporter assays revealed increased 25D1 signals, but not GFP, in the absence of eIF5B (Fig. 6c and Extended Data Fig. 7e).

**Fig. 6.**
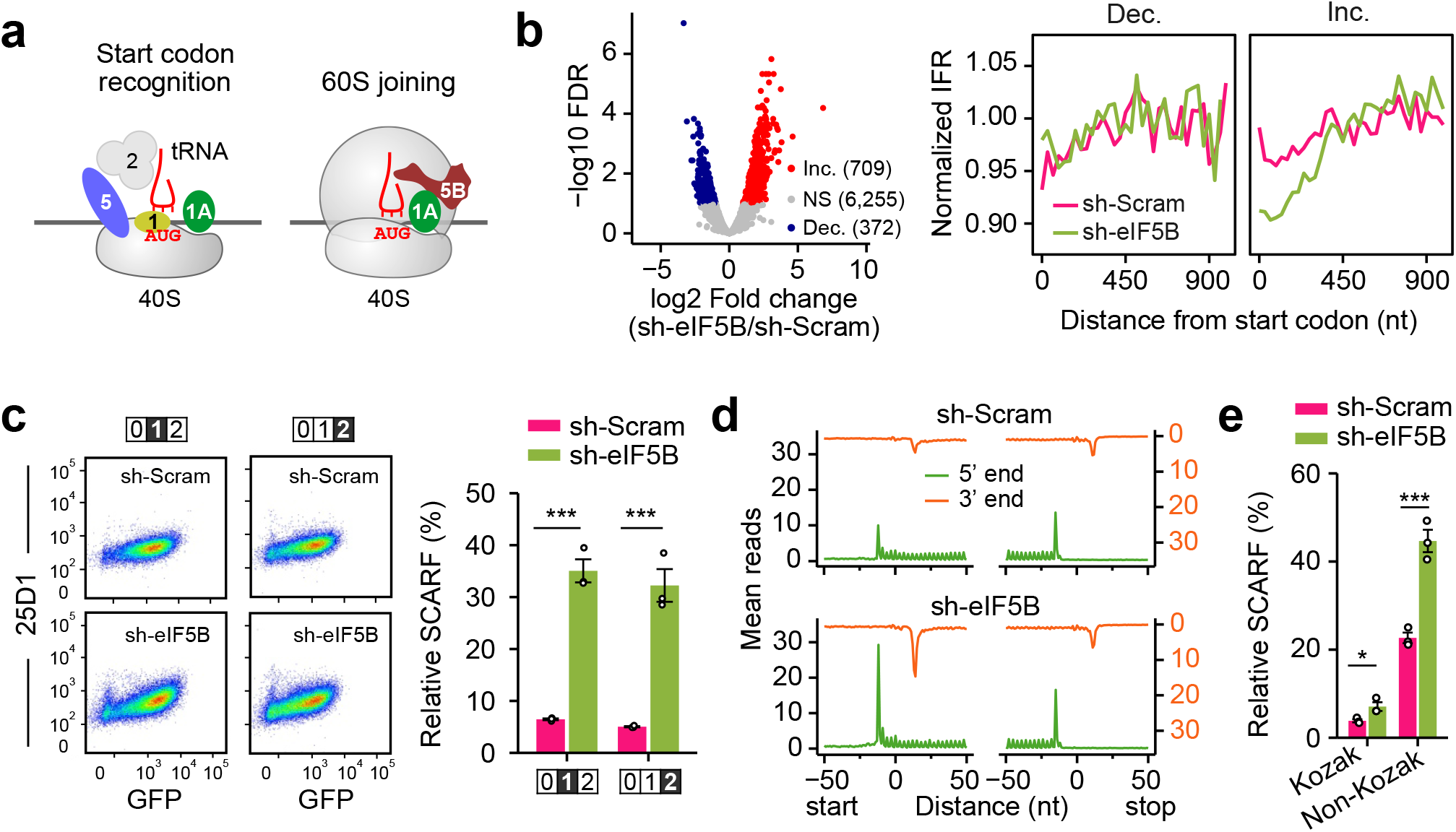
SCARF regulation by eIF5B. **(a)** Schematic of translation initiation factors involving in start codon recognition and 60S subunit joining. **(b)** The left panel is a volcano plot showing the fold change of ribosome density after eIF5B knockdown. The mRNAs with significantly increased ribosome density (FDR < 0.05) are highlighted in red and defined as Inc. The mRNAs with significantly decreased ribosome density (FDR < 0.05) are highlighted in blue and defined as Dec. The right panels show the comparison of IFR along CDS of the two mRNA groups in cells with or without eIF5B knockdown. IFR values are calculated within a non-overlapping sliding window (45 nt), which was subsequently normalized by CDS IFR. **(c)** Representative flow cytometry scatter plots of HEK293-Kb cells transfected with SCARF reporters with or without eIF5B knockdown. Bar plots show the relative SCARF calculated by 25D1 mean fluorescence intensity (MFI) of SCARF reporters over the in-frame control. The highlighted numbers refer to the reading frame of the encoded SIINFEKL. Error bars, mean ± s.e.m.; two-tailed t-test, *n* = 3, *** *P* < 0.001. **(d)** Aggregation plots show the ribosome density across the transcriptome in cells with or without eIF5B knockdown. Both 5’ end (green) and 3’ end (orange) of footprints are used for plotting. **(e)** A bar graph shows the relative SCARF rate in HEK293 cells with or without eIF5B knockdown. The relative SCARF rate calculated by the HiBit activity (Frame 1/Frame 0). Error bars, mean ± s.e.m.; two-tailed *t*-test, *n* = 3, * *P* < 0.05, *** *P* < 0.001.

As a universally conserved initiation factor, eIF5B gates the transition of 80S from initiation to elongation. Indeed, eIF5B depletion from HEK293 cells markedly elevated the ribosome density at the start codon (Fig. 6d). A prior study reported that eIF5B tends to have longer residence time on initiating 80S ribosomes formed at non-Kozak start codons ^8^. Intriguingly, transcripts bearing weak start codons are more susceptible to SCARF in cells lacking eIF5B (Extended Data Fig. 8a). A start codon without the Kozak sequence is often associated with leaky scanning ^18^. To distinguish SCARF from leaky scanning, a new reporter is needed because the SIINFEKL-based reporter is not sensitive enough to measure SCARF from weak start codons. We replaced the tracer peptide SIINFEKL with a nano-luciferase-derived peptide HiBit that can be quantified by luminometry with exquisite sensitivity ^31^ (Extended Data Fig. 8b). By placing HiBit at the reading frame 1, we confirmed that the HiBit signal represented SCARF because mutation of the start codon AUG to ACC eliminated HiBit signals (Extended Data Fig. 8c). To measure the leaky scanning in parallel, we inserted the Fluc sequence further downstream. This design allows us to monitor both SCARF and leaky scanning from the same samples by measuring HiBit and Fluc levels, respectively. As expected, a weak start codon led to an increased leaky scanning by >20 fold (Extended Data Fig. 8d, left panel). Upon normalization to the in-frame HiBit signals, the same reporters showed a 5-fold increase of SCARF when the start codon was switched from strong to weak (from 4% to 21%) (Fig. 6e and Extended Data Fig. 8d, right panel). Therefore, a start codon without the Kozak sequence tends to be skipped, resulting in leaky scanning. But once selected, the assembled 80S ribosome is susceptible to SCARF.

In cells lacking eIF5B, we readily observed increased SCARF as evidenced by the elevated HiBit signals relative to the in-frame translation (Fig. 6e). Notably, eIF5B depletion nearly doubled SCARF when the start codon lacks the Kozak sequence. The unexpected role of eIF5B in maintaining the start codon reading frame is consistent with the unique position of eIF5B within the initiating 80S complex ^30,32^. By stabilizing the initiator tRNA at the P-site (Extended Data Fig. 8e), the presence of eIF5B likely prevents the slippage of the initiating ribosome when the start codon is suboptimal.

### Nutrient stress induces SCARF via eIF5B degradation

Despite the quality control by eIF5B in maintaining the start codon-associated reading frame fidelity, SCARF is seemingly an energy-wasting process by generating degradative polypeptides. Cells under nutrient starvation require intracellular amino acid recycling to support essential protein synthesis ^33^. To examine whether nutrient stress induces SCARF, we applied Ezra-seq to HEK293 cells with or without amino acid deprivation. Remarkably, the IFR in starved cells was substantially reduced in the beginning of CDS (Fig. 7a). This was not due to repressed global protein synthesis because only mRNAs with relatively high ribosome occupancy experienced increased SCARF (Extended Data Fig. 9a). Amino acid starvation-induced SCARF was further confirmed by SIINFEKL-based SCARF reporters (Fig. 7b). HiBit-based SCARF reporters further revealed that mRNAs with weak start codons are more susceptible to SCARF in response to starvation (Fig. 7c and Extended Data Fig. 9b).

**Fig. 7.**
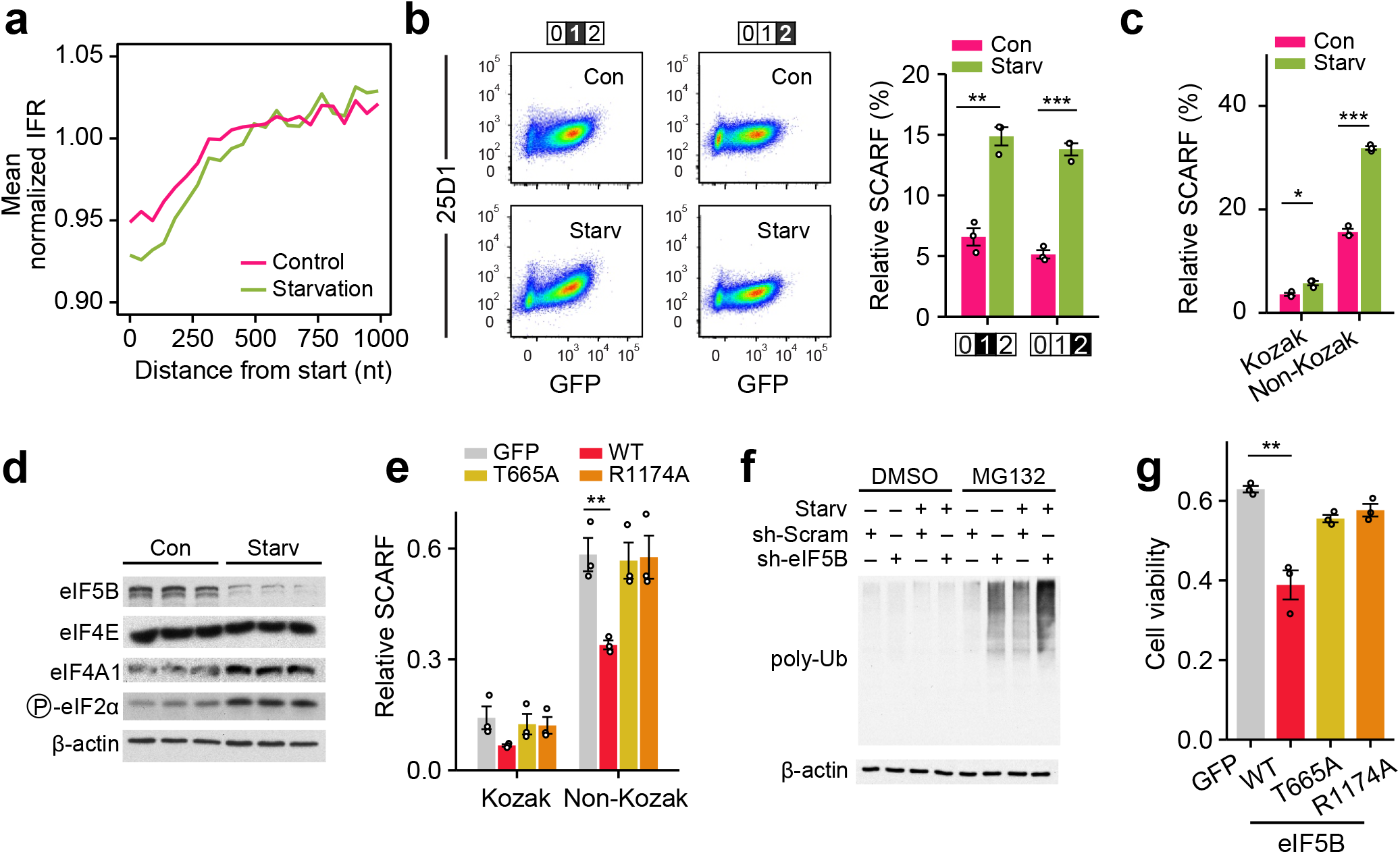
Inducible SCARF by amino acid starvation. **(a)** The comparison of normalized IFR in cells with or without amino acid starvation. IFR values are calculated within a non-overlapping sliding window (45 nt), which was subsequently normalized by CDS IFR. **(b)** Representative flow cytometry scatter plots of HEK293-Kb cells transfected with SCARF reporters before and after amino acid starvation. Bar plots show the relative SCARF calculated by 25D1 mean fluorescence intensity (MFI) of SCARF reporters over the in-frame control. The highlighted numbers refer to the reading frame of the encoded SIINFEKL relative to the AUG codon. Error bars, mean ± s.e.m.; two-tailed t-test, *n* = 3, *** *P* < 0.001. **(c)** A bar graph shows the relative SCARF rate in HEK293 cells after 16 hours amino acid starvation. The SCARF rate was calculated by the HiBit activity (Frame 1/Frame 0). Error bars, mean ± s.e.m.; two-tailed *t*-test, *n* = 3, * *P* < 0.05, *** *P* < 0.001. **(d)** Western blots of translation initiation factors in HEK293 cells before and after 16 hours amino acid starvation. **(e)** A bar graph shows the relative SCARF rate in starved HEK293 cells transfected with GFP, wild type eIF5B, or mutated eIF5B (T665A or R1174A) plasmids. The SCARF rate was calculated by the HiBit activity (Frame 1/Frame 0). Error bars, mean ± s.e.m.; two-tailed *t*-test, *n* = 3, ** *P* <0.01. **(f)** Western blots of polyubiquitinated species in HEK293 cells with or without eIF5B knockdown, in the absence or presence of MG132, before and after amino acid starvation. **(g)** A bar graph shows the cell viability of transfected HEK293 cells after 16 hr of amino acid starvation.

The similar effects on SCARF between eIF5B knockdown and nutrient starvation prompted us to examine whether starvation modulates eIF5B. Intriguingly, amino acid deprivation decreased the steady-state levels of eIF5B, but not other initiation factors like eIF4E or eIF4A1 (Fig. 7d). We found that eIF5B underwent a faster turnover upon nutrient stress (Extended Data Fig. 9c, upper panel) and was stabilized by the proteasome inhibitor MG132 (Extended Data Fig. 9c, bottom panel). Supporting the notion that nutrient stress-induced SCARF is a result of eIF5B degradation, MG132 treatment rescued SCARF by 30% in starved cells (Extended Data Fig. 9d). Additionally, expression of exogenous eIF5B in starved cells suppressed SCARF (Extended Data Fig. 9e and 9f). This result promoted us to conduct rescuing experiments using eIF5B mutants. Recent structural studies revealed a close contact between the domain IV of eIF5B with the acceptor stem of initiator tRNA, whereas the GTPase domain is near the mRNA entry site (Extended Data Fig. 10a) ^32^. We created a G domain mutant (T665A) and a domain IV mutant (R1174A) followed by SCARF measurement in transfected cells. While the wild type eIF5B readily repressed starvation-induced SCARF by 50%, neither the G domain mutant nor the domain IV mutant showed any effects (Fig. 7e and Extended Data Fig. 10b). Therefore, both GTP hydrolysis and interaction with the initiator tRNA are crucial for eIF5B in maintaining the reading frame fidelity during the transition from initiation to elongation.

Nutrient starvation-induced SCARF could supply degradative materials for intracellular amino acid recycling. In line with this notion, either amino acid deprivation or silencing eIF5B resulted in marked accumulation of polyubiquitinated species, which were rapidly degraded (Fig. 7f). In cells subjected to both starvation and eIF5B knockdown, we observed the largest accumulation of polyubiquitinated signals. Additionally, eIF5B overexpression reduced degradative materials in starved cells (Extended Data Fig. 10c). Internal amino acid supply is essential for selective mRNA translation such as ATF4, a cell survival factor during starvation ^34^. Indeed, eIF5B overexpression in starved cells not only suppressed SCARF but also reduced ATF4 expression (Extended Data Fig. 10d and 10e), which was not seen after using eIF5B mutants. We predict that SCARF protects cells from starvation by enabling amino acid recycling and selective mRNA translation. This is indeed the case. Overexpression of wild type eIF5B, but not the G domain mutant (T665A) or domain IV mutant (R1174A), decreased the cell viability during prolonged starvation (Fig. 7g). We conclude that SCARF represents a cellular adaptation mechanism crucial for amino acid homeostasis during nutrient starvation.

## DISCUSSION

Our study demonstrates that start codon recognition does not always guarantee the subsequent reading frame. The unique feature of initiating ribosomes necessitates additional quality control mechanisms to ensure reading frame fidelity, especially when the start codon is suboptimal. Although the Kozak sequence is well-established to facilitate start codon selection, >50% of human transcripts do not contain optimal start codons. The weak start codon not only leads to leaky scanning, but is also susceptible to SCARF, a previously unappreciated phenomenon. SCARF is distinct from alternative initiation because it is coupled to the main start codon. SCARF also differs from elongation-associated frameshifting because it starts with the initiator tRNAi^Met^. As a result, SCARF products are often overlooked from proteomic studies because of out-of-frame translation with non-matched first amino acid. Given the rapid turnover of frameshifted peptides, the contribution of SCARF to the proteome diversity is likely underestimated. The SCARF reporters established here can be readily adapted to distinguish start codon-associated frameshifting from other non-canonical translation events.

The promiscuous translation from the start codon suggests that the transition from initiation to elongation is more complex than we previously thought. The GTPase eIF5B is involved in the correct positioning of the initiator tRNAi^Met^ on the 80S ribosome, thereby controlling the elongation commitment ^8,32^. We propose that eIF5B plays a crucial role in maintaining the reading frame fidelity during the transition from initiation to elongation. Intriguingly, eIF5B levels fluctuate during stress and along different developmental stages ^35,36^. It is likely that eIF5B senses environmental cues, thereby contributing to stress adaptation by increasing translational diversity from existing mRNAs. Nutrient stress rapidly alters the proteome landscape via translational reprogramming ^37^. Upon amino acid deprivation, the general protein synthesis is rapidly suppressed but a subset of mRNAs undergoes selective translation. To support the selective protein synthesis, degradative systems are activated to recycle intracellular amino acids. However, what protein sources are preferentially allocated for degradation remains a debatable subject. Previous studies proposed that a ribosome autophagy (ribophage) pathway supplies internal amino acids during acute nutrient stress ^38^, but systematic quantitation of ribosome inventory showed minimal ribosome degradation ^25^. Given the stress-induced eIF5B degradation, we hypothesize that the subsequently increased SCARF products provide a degradative source for intracellular amino acid recycling during starvation.

A central tenet of biology is the accurate flow of genetic information from nucleic acids to proteins. Biological noises, however, are often overlooked. Translational “noises” derived from SCARF are analogous to divergent transcription at the promoter region by RNA polymerase II ^39^. Exploring translational “noises” under different experimental conditions could indicate whether the translational infidelity arises from error, or represents a potential feature conferring an advantage. For instance, tRNA misacylation and ribosome recoding have been shown to protect cells from oxidative stress ^40^. Based on the nutrient stress inducible feature, the physiological significance of SCARF is twofold: first, it offers an additional means to turn off translation of mRNAs already engaged with ribosomes. Second, it provides an immediate source for intracellular amino acid recycling to enable selective protein synthesis during prolonged starvation. It is equally possible that some of SCARF products could be functional by acting as signaling factors. Broadly, the discovery of SCARF makes sense of translational “noises”, illustrating the beneficial effect of translational diversity in nutrient stress adaptation.

## Supporting information

Supplementary data sets

## Acknowledgements

We thank Cornell University Life Sciences Core Laboratory Center for sequencing and FACS support. This work was supported by US National Institutes of Health (R01GM1222814 and DP1GM142101) and HHMI Faculty Scholar (55108556) to S.-B.Q.

## Author Contributions

Y.M. and S.-B.Q. conceived the study and designed the experiments. Y.M. conducted the majority of data analysis and L.J. performed the majority of experiments. L. D. contributed to the Ezra-seq development. X.E.S. helped with starvation Ribo-seq. S.-B.Q. and Y.M. wrote the manuscript. All authors discussed the results and edited the manuscript.

## Supplementary Information

Extended Data Fig. 1 to Fig. 10

Materials and Methods

References

## METHODS

### Cell lines and reagents

HEK293-K^b^ cells and HEK293 eIF2α (S51D) cells are maintained in Dulbecco’s Modification of Eagle’s Medium (Corning 10-013-CV) with 10% fetal bovine serum (Sigma 12306C). Antibodies used in the immunoblotting are listed below: anti-eIF5B (Proteintech 13527-1-AP), anti-Ubiquitin (P4D1, sc-8017) antibody and anti-β-Actin (Sigma A5441).

### Construction of plasmids and reporters

The full-length coding sequence of firefly luciferase (Fluc) gene was cloned into pcDNA3.1 vector (Invitrogen) to generate Fluc/pcDNA3.1. PCR was performed to generate products of Fluc-based standard frameshifting reporter and HiBit-based SCARF reporters using Fluc/ pcDNA3.1 as a template. PCR products of SIINFEKL-based reporters were generated using a two-step PCR amplification approach. First, the full length SIINFEKL followed by EGFP was amplified from pcDNA3-EGFP to generate SIINFEKL-EGFP. The resulting PCR product was used as a template to produce the full-length reporter using a second forward primer. For exogenous eIF5B, a truncated human eIF5B (580-1220) was cloned into pcDNA3.1(myc-His B) using BamH I and Not I restriction sites. To create the eIF5B mutant, site-directed mutagenesis was performed using Q5 Site-Directed Mutagenesis Kit (New England Biolabs) according to the manufacturer manual. Mutation was confirmed by Sanger DNA sequencing. DNA sequences of all primers used in this study are listed in the Figure S4.

### *In vitro* transcription

To prepare mRNA reporters, 1~2 μg PCR products described above were utilized as templates to generate mRNAs suitable for transfection. *In vitro* transcription was performed with mMESSAGE mMACHINE T7 transcription kit (Ambion) followed by poly(A) tailing kit (Ambion). mRNAs were purified following the manufacturer’s instruction and eluted in nuclease-free water.

### Transfection

For mRNA reporter transfection, cells were transfected with 3 μg mRNA at a mass ratio of 1:1 using Lipofectamine MessengerMAX (Invitrogen) according to the manufacturer’s guide. The mixture was added to cells immediately followed by real time Fluc assay. Alternatively, cells were incubated at 37 °C for 5 h followed by flow cytometry assay or HiBit assay. For eIF5B and eIF5 overexpression, 4 μg plasmids were mixed with 8 μL Lipofectamine 2000 (Invitrogen) and the mixture was added to cells followed by incubation at 37 °C for at least 24 h. For flow cytometry assay, cells were transfected with SIINFEKL-based reporter mRNAs followed by amino acid starvation treatment. For HiBit assay, cells were transfected with HiBiT dual-luciferase mRNA reporter followed by amino acid starvation treatment.

### Firefly luciferase and HiBit assay

Cells grown in 35 mm dishes were transfected with *in vitro* synthesized luciferase reporter mRNAs (3 μg). Then, firefly luciferase substrate d-luciferin (Regis Tech) was added into the culture medium (1 mM) and gently mixed immediately after transfection. Luciferase activity was monitored and recorded using Kronos Dio Luminometer (Atto). HiBit and firefly luciferase activity were measured simultaneously using a Nano-Glo HiBiT Dual-Luciferase Reporter System kit (Promega, N1630) with purified LgBiT Protein (Promega). Briefly, Fluc activity was measured first following addition of the ONE-Glo EX Reagent, which contains detergent to lyse cells and the firefly luciferin substrate. NanoDLR™ Stop & Glo® Reagent and LgBiT protein were then added to quench the Fluc signal and to provide the substrate needed to measure the activity of the HiBiT-LgBiT complexes. HiBit and Fluc activities were recorded using Luminometer.

### Cell treatment

Amino acid starvation treatment was carried out by incubating cells in HBSS buffer (Lonza) with 10% dialyzed FBS (Sigma Aldrich). Samples were collected at indicated time points. For protein turnover and ubiquitination assay, cells were treated with cycloheximide (CHX) at 100 μg/ml or with 5 μM of MG132, followed by collection at indicated time points. HEK293 eIF2α (S51D) cells were treated with 250 nM doxycycline for 48 h for eIF2α(S51D) (DAA-A681) mutant allele.

### Flow cytometry

Transfected HEK293-K^b^ cells were washed with PBS and harvested by trypsin. Cells were pelleted at 2000 rpm for 2 min at 4 °C followed by resuspension in the blocking buffer (1% bovine serum albumin in PBS). Cells were aliquoted into a 96-well plate followed by centrifugation at 2000 rpm for 2 min. Cells were washed one more time followed by staining with 25D1 Alexa 647 antibody (1:1000). After incubation in the dark at 4 °C for 30 minutes with gentle rocking, cells were washed three times with the blocking buffer to remove unbound antibodies. Resuspended cells were subjected to single cell filtering (Falcon) followed by analysis on a BD FACSAria Fusion flow cytometer (BD Biosciences). Analysis of the flow cytometry data was performed using FlowJo.

### Immunoblotting

Cells were washed twice in ice-cold PBS and lysed on ice in SDS-PAGE sample buffer (50 mM Tris pH 6.8, 100 mM dithiothreitol, 2% SDS, 0.1% bromophenol blue, 10% glycerol), followed by heating for 10 min at 95 °C. Proteins were separated on SDS-PAGE and transferred to PVDF membranes (Fisher). Subsequently, membranes were blocked in TBS containing 5% non-fat milk and 0.1 % Tween-20 for 1 h, followed by incubation with primary antibodies overnight at 4 °C. After incubation with horseradish peroxidase-coupled secondary antibodies at room temperature for 1 h, immunoblots were visualized using enhanced chemiluminescence (GE Healthcare).

### shRNA knockdown

shRNA targeting eIF5B was designed from BROAD RNAi consortium database (https://portals.broadinstitute.org/gpp/public/) and listed in Table S1. Oligos were annealed and then cloned into DECIPHER pRSI9-U6-(sh)-UbiC-TagRFP-2A-Puro (Cellecta) according to the manufacturer’s instructions. Lentiviral particles were packaged using Lenti-X 293T cells (Clontech) following the instructions. Virus-containing supernatants were harvested after 48 h transfection and filtered by syringe through a 0.45 μM filter (Millipore) to eliminate cell contaminates. HEK293-K^b^ cells were infected by shRNA lentivirus for 48 h before selection by 2 μg/mL puromycin. Knockdown efficiency was verified by western blotting. A scrambled shRNA was used as control.

### Cell viability assay

HEK293-K^b^ cells were preincubated in a 96-well plate for 24 h in a density of 2000 cells per well. Cell viability was evaluated using the Cell Counting Kit-8 (CCK-8) (APExBIO) according to the manufacturer’s instructions.

### Polysome profiling

A total of 4 plates (10-cm) HEK293-K^b^ cells grown to 80% confluency were changed with fresh medium to remove the dead cells 2 h before harvesting. Cells were washed by cold PBS and lysed in the polysome lysis buffer (10 mM HEPES, pH 7.4, 100 mM KCl, 5 mM MgCl2, 100 μg/mL cycloheximide with 2 % Triton X-100). The lysates were cleared by centrifugation at 14,000 rpm for 10 min at 4 °C. 15-45 % (wt/vol) sucrose density gradients were freshly prepared in a SW41 ultracentrifuge tube (Backman) using a Gradient Master (BioComp Instruments). 500 μL of cytosolic extracts were loaded onto sucrose gradients followed by ultracentrifugation for 2 h 30 min at 32,000 rpm 4 °C in a SW41 rotor. Polysome profiles were recorded at A254 using the Brandel Gradient Fractionation System and an ISCO UA-6 UV/Vis detector. An aliquot of ribosome fractions representing monosome or polysome were collected followed by digestion with *E. coli* RNase I (Ambion, 750 U per 100 A260 units) by incubation at 4 °C for 1 h. Total RNA was extracted using Trizol LS reagent (Invitrogen). Purified RNAs were used for cDNA library construction.

### Ezra-seq and deep sequencing

The ribosome-protected mRNA fragments were separated on a 15% polyacrylamide TBE-urea gel (Invitrogen) and visualized using SYBR Gold (Invitrogen). Selected regions in the gel corresponding to 25–35 nt were excised. RNA fragments were dissolved by soaking overnight in 400 μL RNA elution buffer (300mM NaOAc, pH 5.2, 1mM EDTA, 0.1 U mL^−1^ SUPERase_In). The gel debris was removed using a Spin-X column (Corning), followed by ethanol precipitation. Purified RNA fragments were resuspended in nuclease-free water and quantified using Qubit 2.0 Fluorometer (Invitrogen). A fraction of RNAs (10~200 ng) were used for cDNA library construction. In brief, 10 μL RNAs were mixed thoroughly with 1 μL T4 PNK (10 U, NEB), 1 μL purified TS2126 and 1 μL PAP1 enzyme (NEB) in 7 μL Ezra buffer). The mixture was incubated at 37°C for 30 min followed by 65°C for 20 min. After ethanol precipitation at −20°C for 30 min, the dissolved RNA pellet was mixed with 0.5 μM 5’ end adaptor (Extended Data Table S4) followed by ligation for 90 min at 22 °C in a 10 μL reaction mixture (1 × T4 Rnl2 reaction buffer, 10 U SUPERase_In, 15% PEG8000 and 20 U T4 RNA ligase 2 truncated KQ (NEB)). The ligated RNA sample was mixed with 1 μl 0.5 mM dNTP and 1 μl 0.5 μM RT primer (Extended Data Table S4) and incubated at 65 °C for 3 min, followed by incubation on ice for 1 min. The reaction mix was then added with 9 μl cDNA synthesis mix (5 × first strand buffer, 0.1 M DTT and 750 μg/μl m-MLV-mut5) followed by incubation at 50°C for 30 min. The synthesized cDNAs were mixed with 10 μL rRNA depletion oligo mix tagged with biotin (1 μL 25 μM oligo mix and 3 μL 20 × SSC buffer) and heated at 94°C for 30 s followed by slowly cooling down (3 °C/min) to 37 °C. 30 μL pre-washed streptavidin magnetic beads was added to each sample and stay at room temperature for 10 min. After placing on magnet stand for 1 min, the supernatant was transferred to a new tube and precipitated by ethanol. rRNA-depleted cDNAs were amplified by PCR using barcoded sequencing primers (Extended Data Table S4). The PCR contains 1 × HF buffer, 0.2 mM dNTP, 0.5 μM oligonucleotide primers, and 0.5 U Phusion polymerase. PCR was carried out under the following conditions: 98 °C, 30 s; (98 °C, 5 s; 68 °C, 15 s; 72 °C, 10 s) for 12 cycles; 72 °C, 3 min. PCR products were separated on a 8% polyacrylamide TBE gel (Invitrogen). Expected DNA at 180 bp was excised and recovered from DNA gel elution. After quantification by Agilent BioAnalyzer DNA 1000 assay, equal amounts of barcoded samples were pooled into sequencing lanes, which were sequenced using NextSeq 500 (Illumina).

## DATA PROCESSING AND ANALYSIS

### Ribo-seq analysis

#### Sequencing reads alignment

To compile human and mouse transcriptome, we downloaded annotation files from ENSEMBL database (GRCh38.81 for human, and GRCm38.83). Protein coding transcripts were extracted based on the annotation files, using house-hold scripts. For each gene, the transcript with longest CDS was selected. In the case of equal CDS length, the longest transcript was used. rRNA sequences were downloaded from the nucleotide database of NCBI and RNAcentral^1^. To align sequencing reads, the 3’ adapters of the reads were trimmed by Cutadapt ^2^, using parameters: -a AAAAAA --max-n=0.1 -m 15. The trimmed reads with length shorter than 15 nucleotides (nt) were excluded from analysis. To keep accurate reading frame, low-quality bases at both ends of the reads were not subject to clip. The clean reads were first aligned to rRNAs using Bowtie^3^, with parameters: -v0 --norc. The unaligned reads were then mapped to human or mouse transcriptome using STAR^4^ with default parameters. To avoid ambiguity, reads mapped to multiple positions or with > 2 mismatches were disregarded for further analysis. Reads with unaligned bases at the 5’ end (i.e. flagged as soft-clip by STAR aligner) were further excluded from analysis. Ribosome P-site was defined as the positions of 12^th^, 13^th^ and 14^th^ from 5’ end of the read (position 0). A-site was defined as the positions of 15^th^, 16^th^ and 17^th^.

#### Mean footprint density along CDS

For each mRNA, footprints at individual sites were normalized by mean footprints of the CDS. mRNAs with total reads in CDS < 16 or the CDS sites covered by footprints < 10% were excluded. The normalized values of the sites with the same distance relative to the start codon or stop codon were averaged across transcriptome.

#### In-frame ratio along CDS

We first calculated codon in-frame ratio. For each codon position within a CDS, the in-frame ratio was calculated by dividing in-frame footprint by the total footprints at the codon. The codons with total footprints < 10 were excluded. Then, in-frame ratios of the codons at the same distance relative to start or stop codons were averaged across transcriptome. A bootstrap method was performed to estimate the variation of mean in-frame ratio. To this end, we repeated the process to calculate the mean in-frame ratios along CDS, by using an artificial transcriptome, which was generated by randomly picking up mRNAs from real transcriptome. The artificial transcriptome has equal number of mRNAs to the real transcriptome. The standard variation was calculated based on the mean in-frame ratios from 100 artificial transcriptomes. To compare mean in-frame ratios between mRNA groups or samples, we instead calculated in-frame ratio using a non-overlapping sliding window with 30 nucleotides in length. First, same as codon inframe ratio, in-frame ratio of the window was calculated. Then, mean in-frame ratios were calculated by average in-frame ratios of the windows located at the same distance to start or stop codons. Because both starvation and eIF5B knockdown obviously inhibit global translation, the sliding window was increased to 45 nucleotides to reduce the noises of in-frame ratio.

#### In-frame ratio at ribosome A-site or P-site

Total footprints at the 61 codons (not including the three stop codons) were counts respectively, based on the reads with A-site or P-site located within CDS. To focus our analysis on elongation, the first and last 30 codons of CDS were excluded from analysis. For each codon, read ratios in different reading frames were calculated as the ratio of the reads with P-site or A-site located to in-frame, frame 1 or frame 2 respectively over the total reads at the codon. To study the relationship of codon optimality to reading fidelity, the 61 codons were divided into optimal and non-optimal codons, based on tRNA adaptation index (tAI) ^5^. Codons with tAI < 0.3 were defined as non-optimal codons, and the others were optimal codons.

#### Codon occupancy at ribosome A-site or P-site

For each mRNA, footprints at the same codons within CDS were counted, and then normalized by the average footprint of the CDS. mRNAs with CDS footprints < 32 were not included. Footprints with P-site or A-site located to the first and last 30 codons were also excluded. The normalized ribosome occupancies at the same codons were averaged over the transcriptome.

#### Differential genes upon eIF5B knockdown

Ribosome footprints mapped to CDS of individual mRNAs were counted using a house-hold script. mRNAs with total reads in all samples < 10 were excluded. The count table was then analyzed by DESeq2 ^6^. mRNAs with false discovery rate (FDR) < 0.05 and fold change > 2 were defined as significantly up-regulated mRNAs. mRNAs with FDR < 0.05 and fold change < 0.5 were significantly down-regulated mRNAs. The significantly changed mRNAs were then analyzed by PANTHER^7^ to search enriched biological processes and pathways.

#### Calculation of initiation potential

We used ribosome footprints in 5’ UTR to calculate the initiation potential of a triplet when a scanning subunit reaches the triplet. If a triplet can serve as a start codon, the reads in downstream coding frame relative to the triplet should become higher than the reads within the same frame before the triplet. With this assumption, for all triplets in 5’ UTR, we counted IFR values in a small window (30 nt or to a stop codon if the ORF is shorter than 30 nt) after the triplet, which was then subtracted from the IFR value in the window (30 nt) before the triplet. The triplets with total reads in any of the two windows <5 were not included. Considering that AUG usually serve as a strong initiation codon, we also excluded the triples containing AUG in any of the two windows. Although multiple alternative initiation sites in 5’ UTR may overlap each other, thereby affecting the IFR difference between the two windows, an average of IFR difference values across the same triplets in 5’ UTR can eliminate the effect from overlapping hidden ORFs and other random noisy, thus providing an overview of initiation potential of a triplet. For a triplet, we defined the average IFR difference as the initiation potential of the triplet.

#### uORF prediction

We predict uORFs with robust translation based on Ezra-seq data. For each mRNA, we first extracted all possible uORFs, i.e., mRNA regions starting with ATG or the other 10 non-canonical initiators (as indicated by a higher initiation potential), and ending with TAG, TGA or TAA. We used the method from ^8^ to find the potential uORFs. In brief, we applied a Wilcoxon test to test whether the in-frame reads are significantly higher than the other two frames. The two *P* values were then combined to a single *P* value using a Stouffer’s method. The *P* values were then corrected by a false discovery rate (FDR) method. The uORFs with FDR <0.1 were defined as the uORFs with robust translation. The overlapping uORFs located to the same coding frame were merged into a single one, by selecting the stop codon located at the most 5’ end of transcript, and the start codon which maximizes the fraction of in-frame reads.

### mRNA sequence analysis

#### Definition of mRNAs enriched with non-optimal codons

To compare in-frame ratio between mRNAs groups with different fractions of non-optimal codons in the beginning of CDS, we calculated a geometric mean of tAI value for the first 100 codons. mRNAs were categories into four groups based on the quantiles (25%, 50% and 75%) of the mean tAI values.

#### Calculation of Kozak score

We calculated a Kozak score for an mRNA based on a position weight matrix. In brief, we downloaded the initiation strength from previous work ^9^, which evaluated the initiation efficiency by placing random sequences around the AUG codon (i.e., NNNNNNAUGNN). The 100 random sequences with highest initiation efficiency were used to construct a position probability matrix. The position weight matrix (PWM) was calculated using following equation.

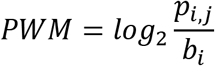

Where, *p_i,j_* refers to the possibility of *i^th^* nucleotide at position *j. i* is an element of (A, T, G, C). *b_i_* is the background frequency of the *i^th^* nucleotides. Here, an equal frequency (i.e., 1/4) was used for simplification.

For each mRNA, we extracted six nucleotides before the start codon, and two nucleotides after the start codon. A Kozak score was defined as the sum of the values of corresponding nucleotides at each position in the PWM. mRNAs were categories into four groups based on the quantiles (25%, 50% and 75%) of the Kozak score.

#### RNA secondary structure analysis

First, a 50 nucleotides sequence with the start codon in the center was extracted The minimum fold free energy (MFE) was calculated by ViennaRNA ^10^ using default parameters. The mRNAs with lowest MFE (bottom 25%) were grouped as start codon structured mRNA, and mRNAs with highest MFE (top 20%) were defined as non-structured mRNA.

### Analysis of massively paralleled reporter assay

#### Count of random sequences

For each sequencing reads, from the raw sequencing file, the first 21 nucleotides at 5’ end, 3’ adapter and low-quality bases were trimmed using Cutadapt. The trimmed reads with length unequal to 10 nucleotides were excluded from analysis. The remaining trimmed reads were counted, and then an RPM value (reads per million) was obtained by dividing the resultant read count by the total count.

#### Triplet frequency along random sequences

We separated mRNA reporters into different groups based on the number of associated ribosomes. For mRNA reporters with sequencing reads > 5, a M/P ratio was calculated as the ratio of RPM value in monosome fraction over RPM value in polysome fraction. The mRNA reporters with the highest M/P ratio (top 10% of the total mRNA reporters) were defined as monosome enriched mRNA reporters. For mRNA reporters sorted by 25D1 and GFP signals, the 25D1 or GFP signals were calculated using the method in Jia et al^11^. The mRNA reporters with the highest ratio of 25D1 signal over GFP intensity (top 10% of the total mRNA reporters) were defined as 25D1-enriched mRNA reporters. We counted triplet frequencies along random sequences inserted in monosome or 25D1 enriched mRNA reporters, respectively.

#### Analysis of variance (ANOVA)

To understand the relative contribution of position-specific nucleotide to frameshifting rate, we conducted a multiway ANOVA^12,13^ using following equation.

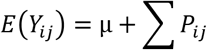

where *Y_ij_* refers to the log transformed ratio of RPM in monosome fraction to RPM in polysome fraction (log2 M/P). *μ* is the overall average effect for all levels. *P_ij_* represents additional effect of the *i^th^* nucleotides at position *j. i* is an element of (A, T, G, C). The combination effects among positions were not considered. We performed ANOVA described above using the “aov” function in R software. The sum-of-squares for each position was directly retrieved from the ANOVA result, accessed using the “summary” function in R. The relative contribution of each position to frameshifting rate was defined as the fraction of sum-of-squares for the position divided by the total sum-of-squares.

### Proteomic analysis

We downloaded human high-quality unidentified spectra from PRIDE cluster database^14^. The searching database includes all proteins encoded by human protein-coding mRNAs, and the predicted peptide sequences generated by out-frame translation initiated from the start codon. For each out-frame peptide, an artificial methionine was added at the N-termini. We also added contaminants obtained from MaxQuant^15^. The searches were performed by X!Tandem^16^ with default parameters. The trypsin was set as the enzyme to digest peptides. The output of X!Tandem was further analyzed by PeptideProphet following the tutorial of Trans-Proteomic Pipeline^17^. TMT-MS spectra were downloaded from An et al^18^. The spectra were analyzed by MaxQuant. For group-specific parameters, the spectra type was set as 11plexTMT for MS3. Oxidation of methionine and N-terminal acetylation were set as variable modifications. Cysteine Carbamidomethylation was set as a fixed modification. The digestion enzyme was set as Trypsin/P and LysC. MS dataset encompassing N-terminal peptides enriched by TAILS was downloaded from Na et al^19^. Acetylation and demethylation of peptide N termini and oxidation of methionine were set as variable modifications. Cysteine Carbamidomethyl modification and dimethyl modification of lysine were set as fixed modifications. The digestion enzyme was set as ArgC. For PRIDE unidentified spectra, all identified peptides were used for metagene analysis. For single mass spectrometry dataset, the peptides with reported FDR < 0.01 and the posterior error probabilities < 0.05 were used. The found peptides from all three types of datasets were exported to Microsoft Excel for further analysis.

### Statistics and reproducibility

For Ezra-seq libraries, two biologically independent replicates were used in most experiment. Immunoblots and flow cytometry plots are the representative images from at least three rounds of independent experiments. Data are presented as the mean ± standard error (s.e.m), with two-tailed student’s *t*-tests or Wilcoxon signed-rank test on the statistical significance of differences between groups. All *P* values have shown in figure legends. All statistical analysis and data graphing were done in R (4.03) software.

