## Supplementary data sets for "Start codon-associated ribosomal frameshifting mediates nutrient stress adaptation"

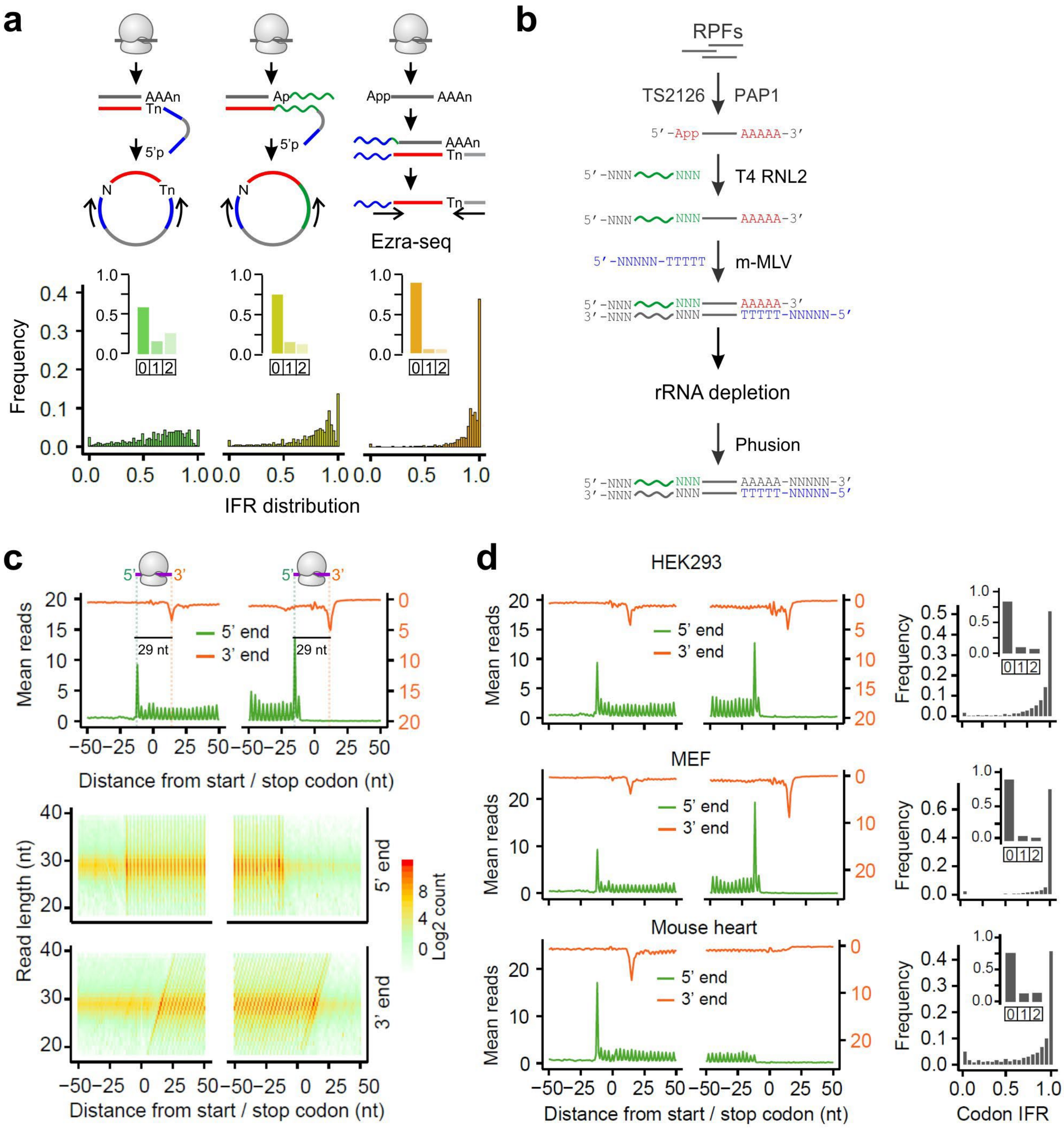


**Extended Data Fig. 1. Development of Ezra-seq.**

**(a)** The top panels show the comparison of Ezra-seq and two ligation-based Ribo-seq procedures. The bottom panels show in-frame ratio (IFR) of reads at codons with 1.0 indicating complete in-frame. The inserted bar plots show the fraction of reads in different reading frames.

**(b)** An outline of Ezra-seq procedure (see Materials and Methods for the detail).

**(c)** Aggregation plots show the ribosome density across the transcriptome. Transcripts are aligned to start and stop codons, respectively. Both 5’ end (green) and 3’ end (orange) of footprints are used for plotting. The heatmap shows the density of ribosome footprints with different length.

**(d)** Aggregation plots show the ribosome density across the transcriptome in different cell lines and tissues. Transcripts are aligned to start and stop codons, respectively. Both 5’ end (green) are 3’ end (orange) of footprints are used for plotting. The right panel shows the range of codon IFR with 1.0 indicating complete in-frame reads. The inserted bar plot shows the fraction of reads in different reading frames.

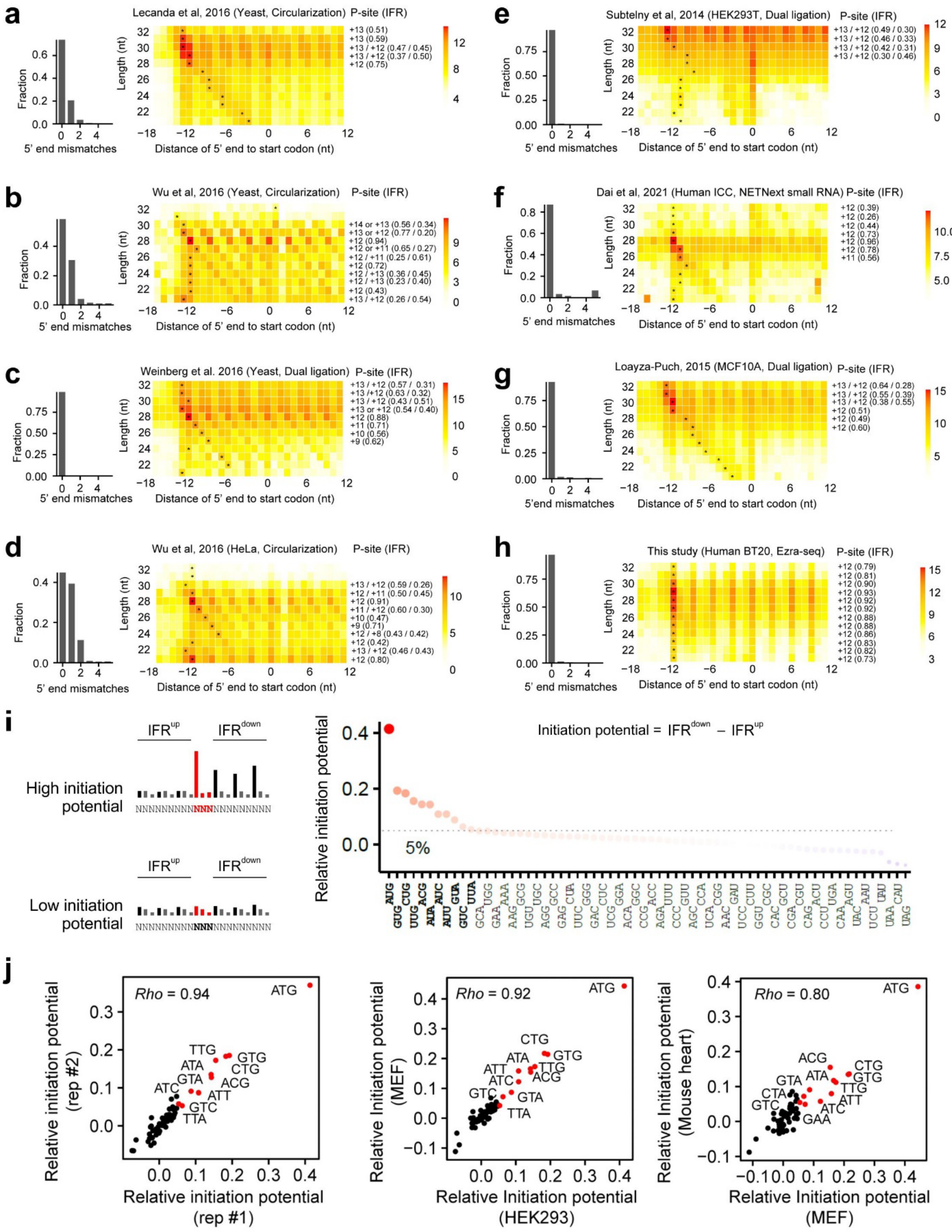

a
Lecanda et al, 2016 (Yeast, Circularization) P-site (IFR)
b
Wu et al, 2016 (Yeast, Circularization) P-site (IFR)
c
Weinberg et al. 2016 (Yeast, Dual ligation) P-site (IFR)
d
Wu et al, 2016 (HeLa, Circularization) P-site (IFR)
e
Subtelny et al, 2014 (HEK293T, Dual ligation) P-site (IFR)
f
Dai et al, 2021 (Human ICC, NETNext small RNA) P-site (IFR)
g
Loayza-Puch, 2015 (MCF10A, Dual ligation) P-site (IFR)
h
This study (Human BT20, Ezra-seq) P-site (IFR)
Fraction
5' end mismatches
Length (nt)
Distance of 5' end to start codon (nt)
i
High initiation potential
Low initiation potential
Initiation potential = IFR^down − IFR^up
Relative initiation potential
5%
j
Rho = 0.94
Rho = 0.92
Rho = 0.80
Relative initiation potential (rep #2)
Relative initiation potential (rep #1)
Relative Initiation potential (MEF)
Relative Initiation potential (HEK293)
Relative Initiation potential (Mouse heart)

**Extended Data Fig. 2. Comparison of 5' end accuracy across representative Ribo-seq data sets.**

**(a)-(h)** For each panel, the left bar plots show the number of mismatches at the 5' end of footprints. The heatmap shows the distance of 5' end to the start codon. All footprints were classified into different length (y-axis), the color represents the $\log_2$ count of footprint. For each length group, the submit peak was indicated by a star, which indicates the distance of P-site to the start codon (the number at the right side of heatmap). The number in parentheses is in-frame rate when the left P-site offset was used.

**(i)** Schematic of IFR changes after a triplet with high or low initiation potential (left panel). A scatter plot shows initiation potential of 64 triplets based on IFR changes (right panel).

**(j)** Correlation of initiation potential of 64 triplets between biological replicates (left), between HEK293 and MEF cell lines (middle), or between MEF cell lines and mouse heart (right). The AUG and 10 non-AUG triplets with highest initiation potential in HEK293 were highlighted in red.

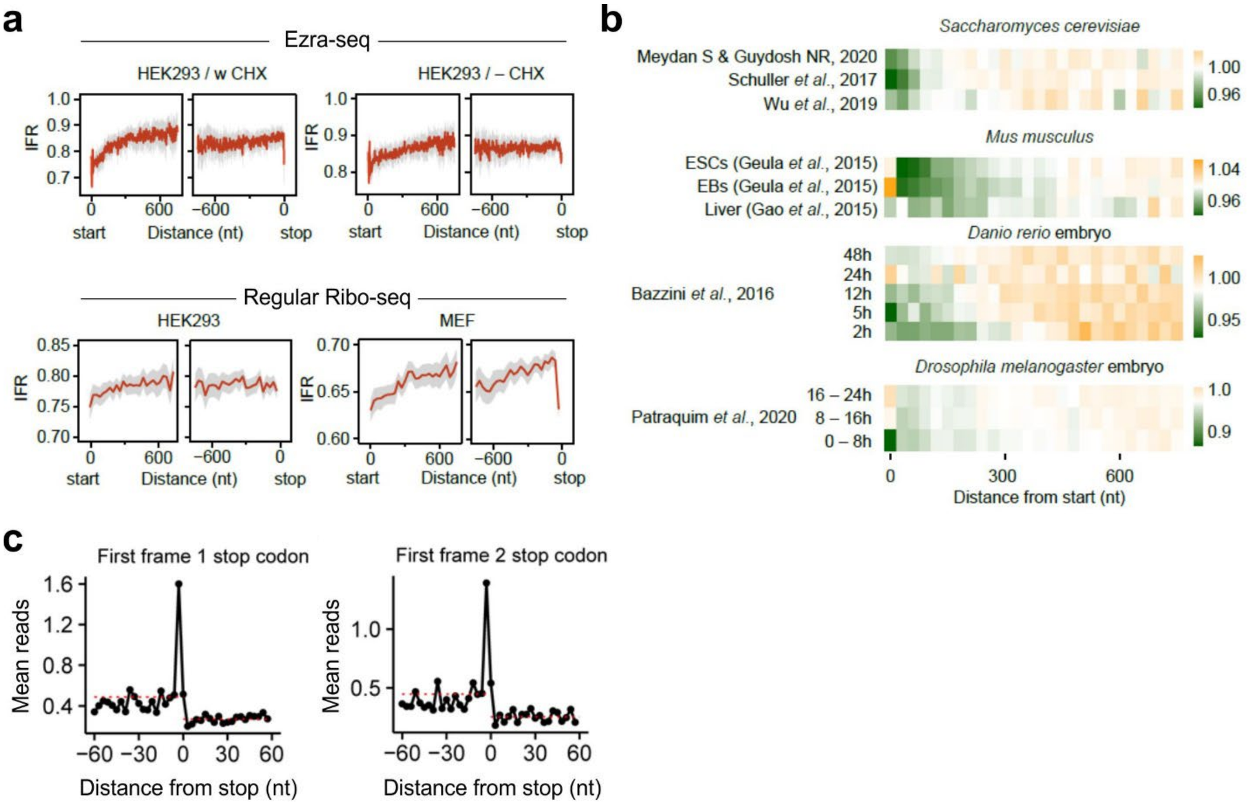


**Extended Data Fig. 3. Characterizing in-frame and out-of-frame RPFs.**

**(a)** Top panel shows in-frame ratio of ribosome footprints across the transcriptome in cells with (left panel) or without (right panel) cycloheximide (CHX) treatment (100 µg/mL) for 30 min. Bottom panel shows in-frame ratio of ribosome footprints across the transcriptome in HEK293 and MEF cells. Ribo-seq data were obtained by ligation-based Ribo-seq methods. Due to the relatively low resolution, IFR values were calculated within a non-overlapping sliding window (30 nt). Grey shadow shows the variation of mean IFR estimated by bootstrap method. Transcripts are aligned to start and stop codons, respectively

**(b)** Heat maps show normalized IFR of CDS across different species and cell lines using published Ribo-seq data sets. IFR values were calculated within a non-overlapping sliding window (30 nt), which was subsequently normalized by CDS IFR.

**(c)** An aggregation plot shows out-of-frame reads around the 1st frame 1 or frame 2 stop codons. The mean out-of-frame reads before and after the out-of-frame stop codons are indicated by dashed lines.

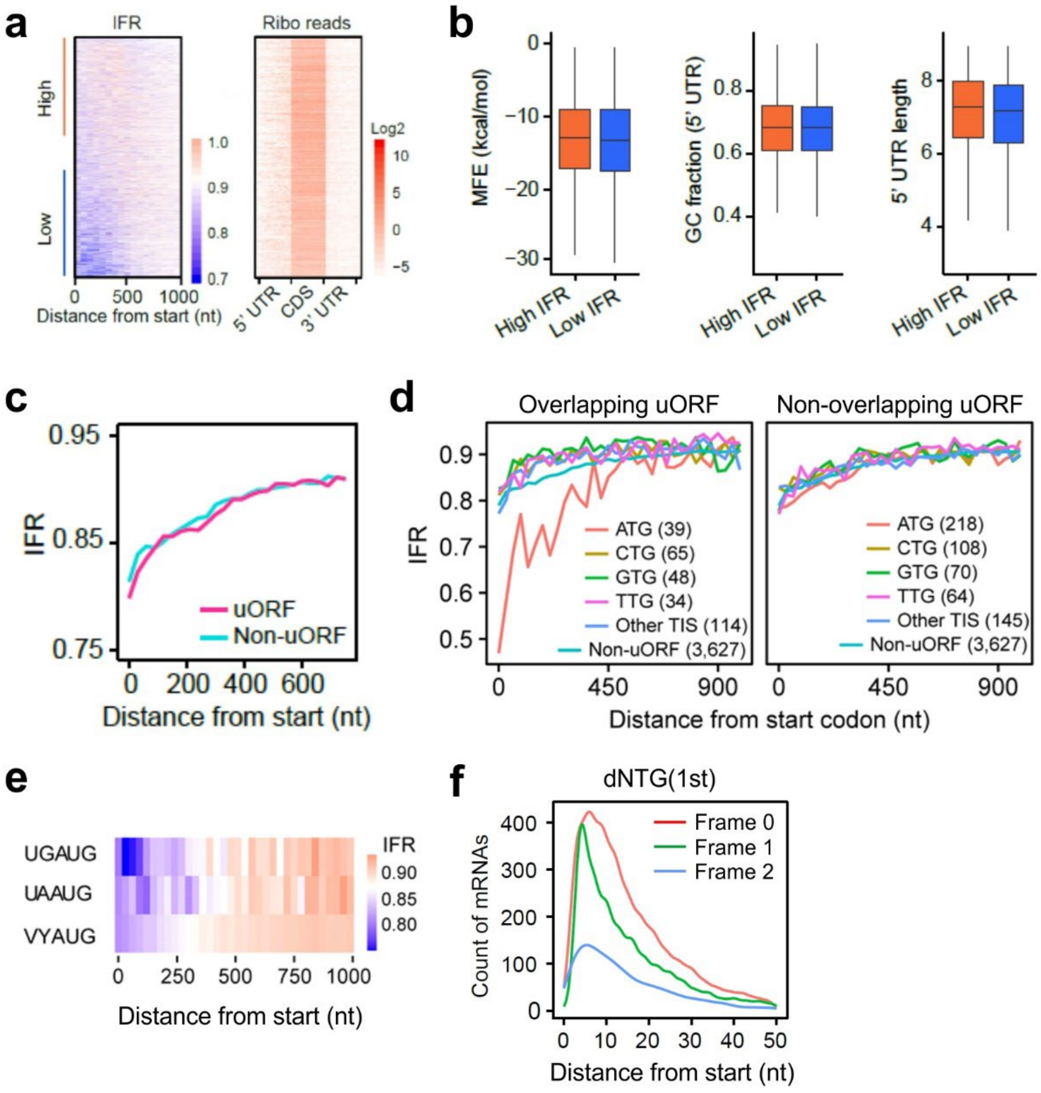


**Extended Data Fig. 4. uORF and leaky scanning minimally contribute to reduced IFR.**

**(a)** A heat map shows the IFR values at the first 333 codons of individual mRNAs. The right heat map shows ribosome densities in different regions of individual mRNAs.

**(b)** Boxplots show RNA fold free energy (MFE) around start codon, GC fraction in 5' UTR and 5' UTR length between mRNAs with low or high IFR in the beginning of CDS.

**(c)** Comparison of IFR between mRNAs with or without uORF. IFR values are calculated within a non-overlapping sliding window (30 nt).

**(d)** Effects on uORF translation on in-frame ratio in the beginning of CDS. uORFs were identified by Ezra-seq data in this study. All uORFs were separated into different groups based on the initiators. Overlapping uORFs are defined if the stop codon of uORF is beyond the start codon of main CDS. The numbers indicate the number of mRNAs used for analysis.

Of note, when uORFs strongly inhibit main CDS translation, those mRNAs were not included in analysis due to the lack of sufficient reads on the main CDS.

**(e)** A heatmap shows IFR values between mRNAs with or without a stop codon UGA before start codon. V: not uracil, Y: pyrimidine.

**(f)** Distance of the first downstream NTG (dNTG) in different reading frames relative to the annotated start codon.

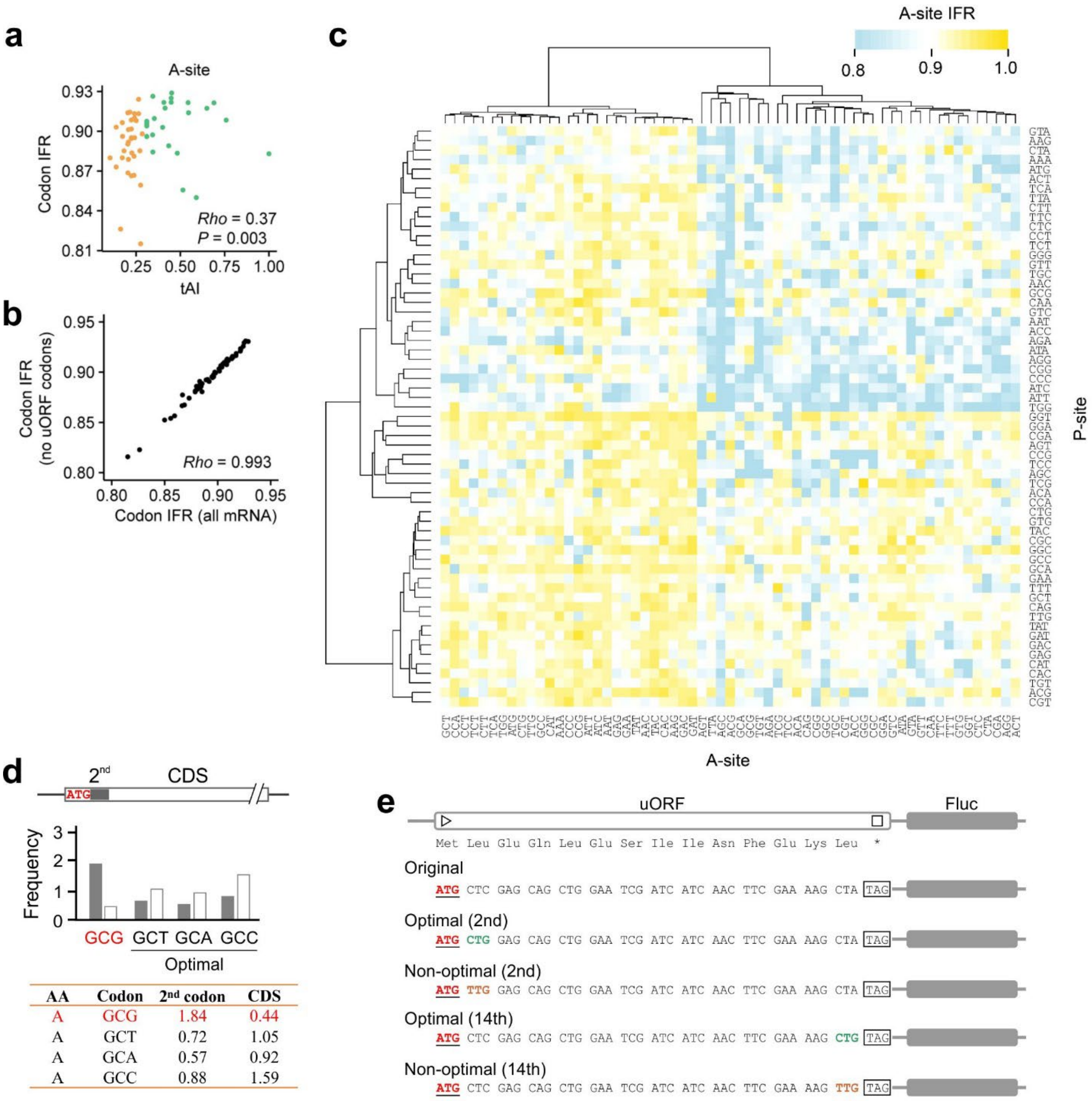


**Extended Data Fig. 5. Non-optimal codons induce ribosome frameshifting.**

**(a)** A scatter plot shows the correlation between codon optimality and IFR when the A-site codon is considered.

**(b)** A scatter plot shows the correlation of IFR values for transcripts with or without uORFs.

**(c)** A heat map shows the effect of P-site and A-site combinations on reading frame fidelity at ribosome A-sites.

**(d)** Analysis of codon usage bias of the first codon after the start codon. Bar plot and the table show the relative synonymous codon usage (RSCU) of the most prevalence amino acid Alanine at the second codon of CDS.

**(e)** Sequences of uORF reporter with either the 2$^{nd}$ codon or 14$^{th}$ codon replaced by a synonymous optimal (green) or non-optimal (orange).

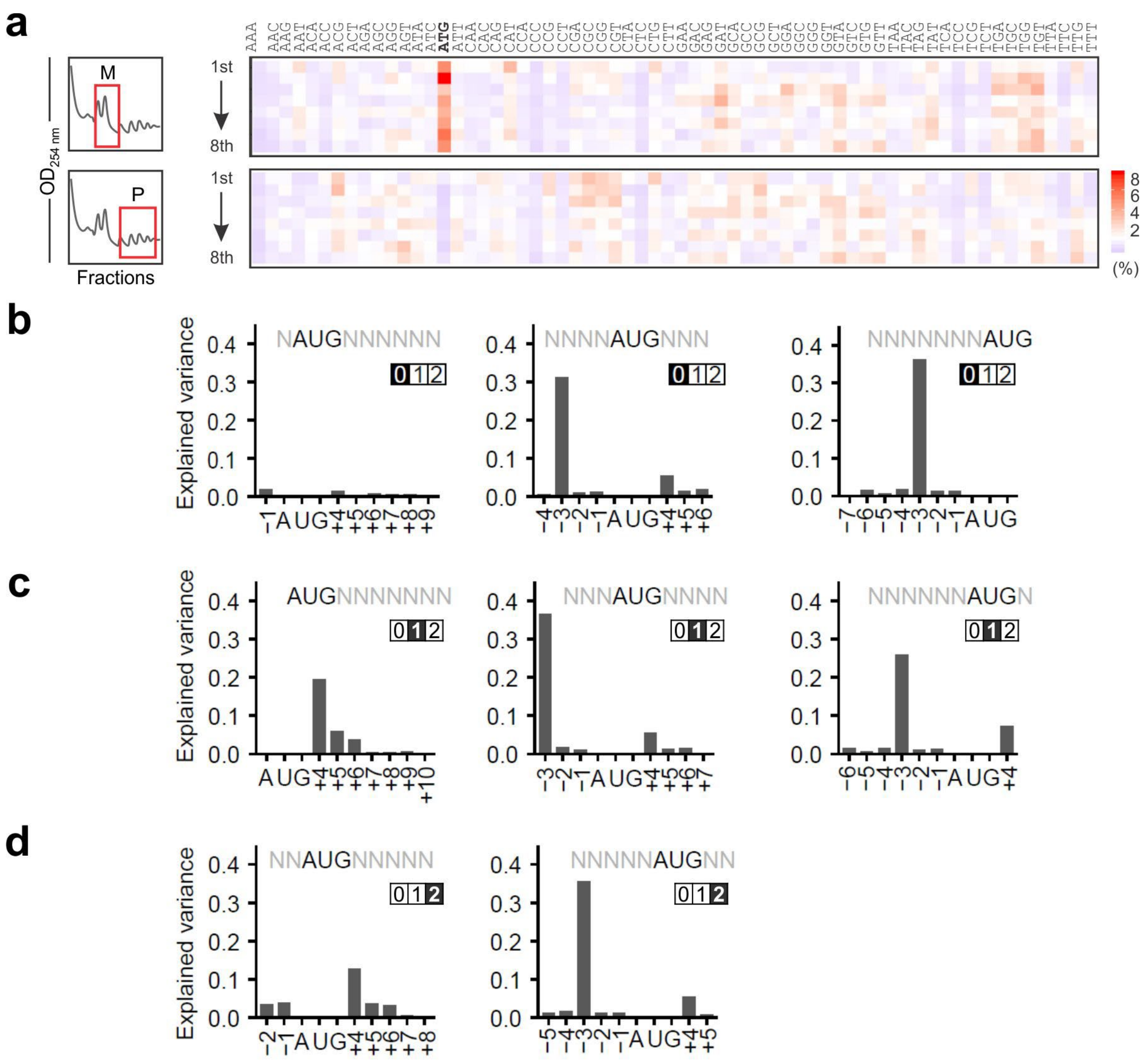

**Extended Data Fig. 6. SCARF regulation by the start codon sequence context.**

**(a)** HEK293-K$^{b}$ cells were transfected with massively parallel mRNA reporters followed by sucrose gradient separation into monosome (M) and polysome (P) fractions. The original frequency of triplets in different populations is shown as heat maps.

**(b – d)** Relative contributions of the nucleotide identity in different positions to the uORF translation based on the M/P ratio. The highlighted numbers refer to the reading frame of the encoded SIINFEKL relative to the AUG codon.

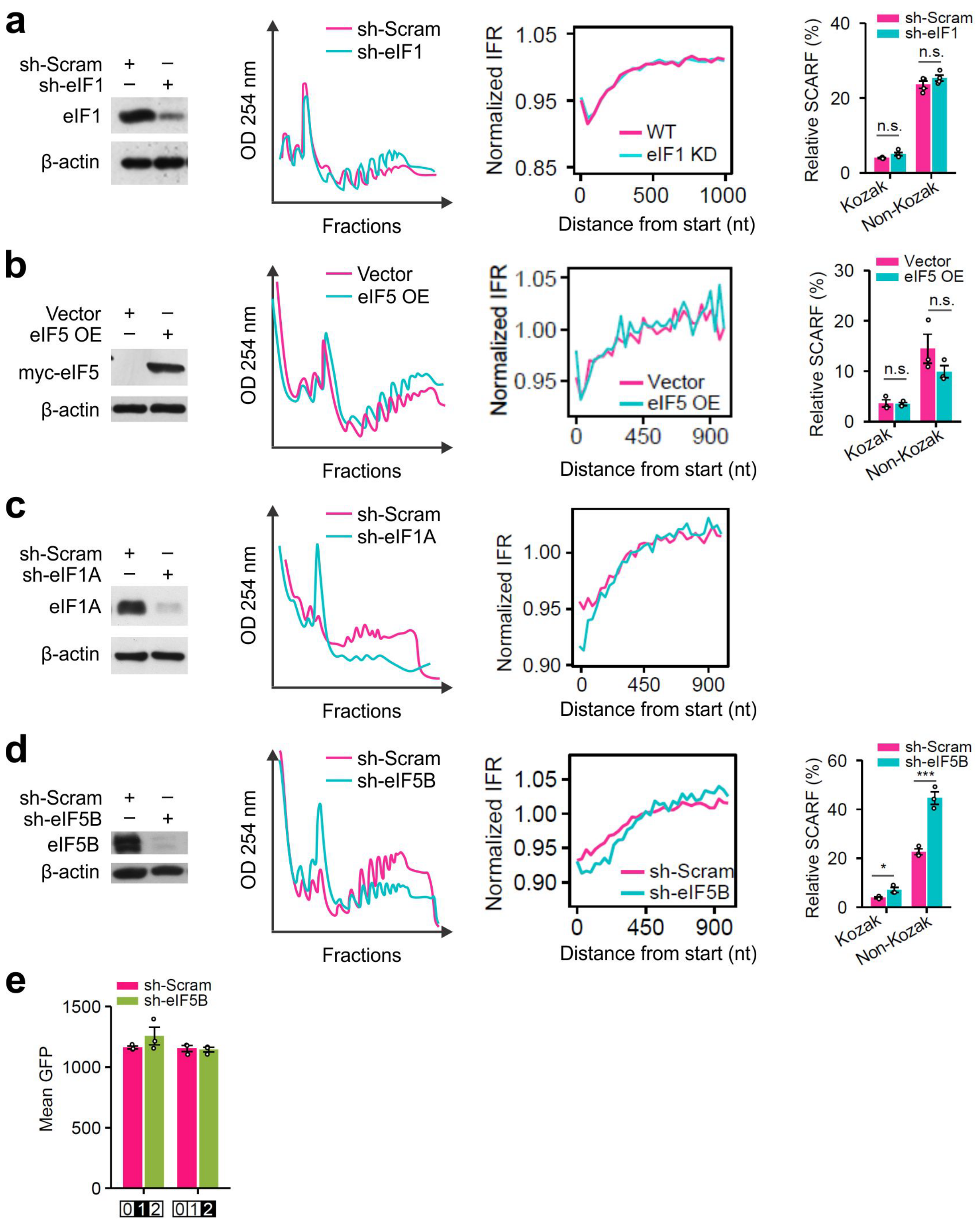


**Extended Data Fig. 7. Characterizing the regulatory role of translation initiation factors in SCARF.**

**(a)** The left panel shows the western blots of HEK293 cells with or without eIF1 knockdown. The middle panel shows the comparison of polysome profiles of cells with or without eIF1 knockdown. The right panel shows the comparison of normalized IFR in cells with or without eIF1 knockdown. IFR values are calculated within a non-overlapping sliding window (45 nt), which was subsequently normalized by CDS IFR. The right panel shows the HiBit-based SCARF reporter assay. Error bars, mean ± s.e.m. n = 3. Two-tailed *t*-test, $n = 3$, n.s. no significant change.

**(b)** Same as (A) using cells with eIF5 overexpression.

**(c)** Same as (A) using cells with eIF1A knockdown.

**(d)** Same as (A) using cells with eIF5B knockdown. Two-tailed *t*-test, $n = 3$, * $P < 0.05$, *** $P < 0.001$.

**(e)** Bar plots show the relative GFP mean fluorescence intensity (MFI) of SCARF reporters over the in-frame control. Error bars, mean ± s.e.m..

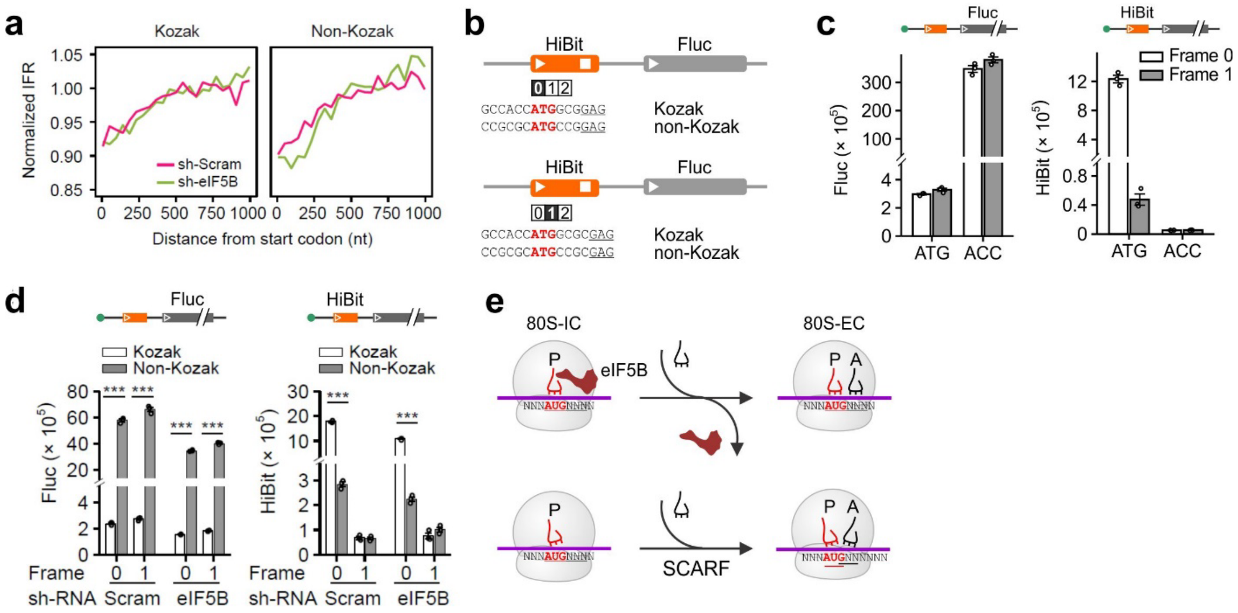


**Extended Data Fig. 8. Characterizing the regulatory role of translation initiation factors in SCARF.**

**(a)** Comparison of normalized IFR for mRNAs with or without the Kozak sequence context on start codons in cells with or without eIF5B knockdown. IFR values are calculated within a non-overlapping sliding window (45 nt), which was subsequently normalized by CDS IFR.

**(b)** Sequence information of HiBit-based SCARF reporter.

**(c)** Bar graphs show the HiBit-based SCARF reporter assays in in HEK293 cells. Both the Fluc (left panel) and HiBit (right panel) signals were measured from cells transfected with SCARF reporters bearing a uORF start codon or a uORF with start codon mutated to ACC. Error bars, mean ± s.e.m.

**(d)** Bar graphs show the HiBit-based SCARF reporter assays in cells with or without eIF5B knockdown. Both the Fluc (left panel) and HiBit (right panel) signals were measured from cells transfected with SCARF reporters bearing a uORF start codon with or without the Kozak sequence context. Error bars, mean ± s.e.m.; two-tailed $t$-test, $n = 3$, *** $P < 0.001$.

**(e)** A model depicting the role of eIF5B in stabilizing initiator tRNA at the P-site, thereby maintaining the reading frame during the transition from initiation to elongation.

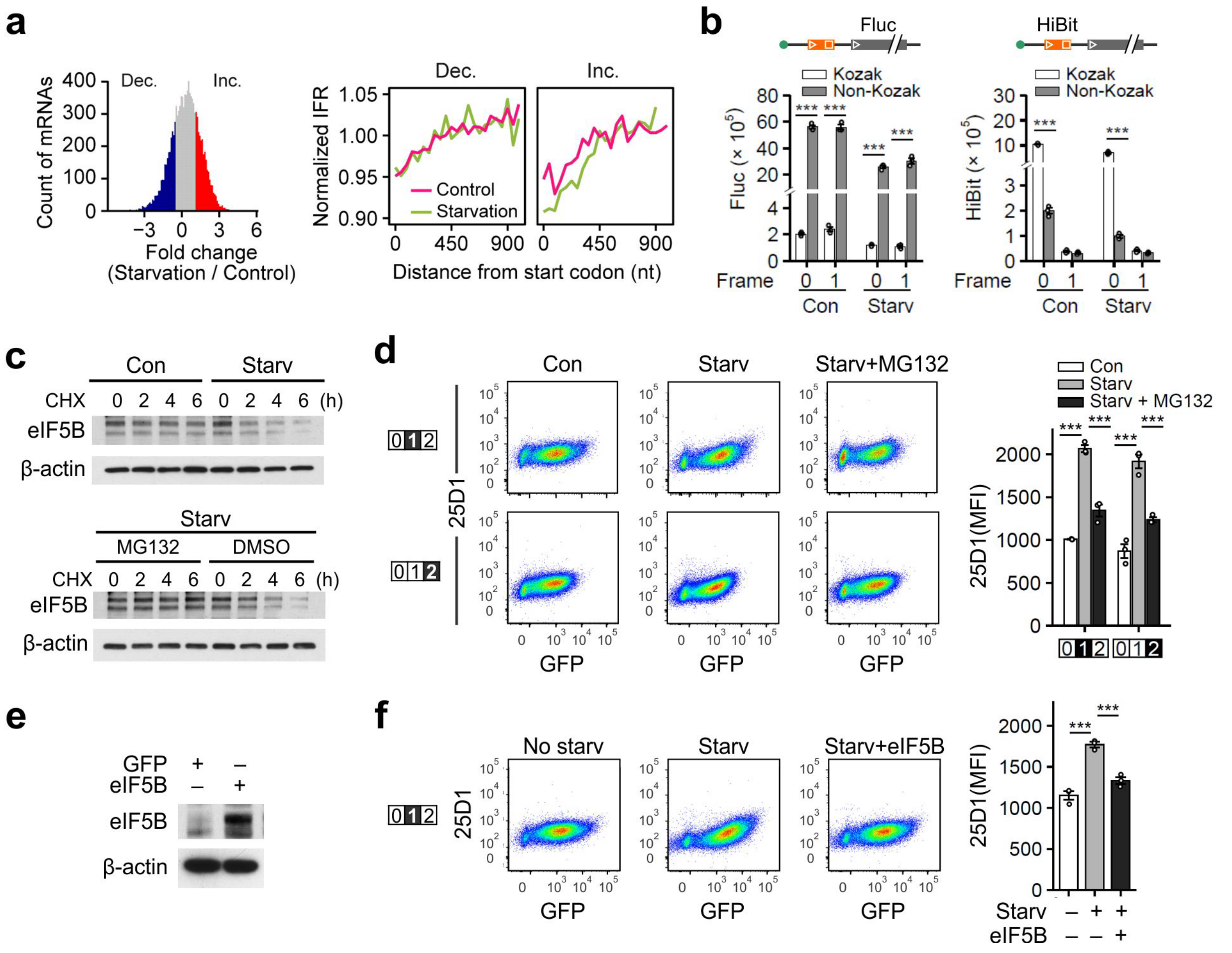


**Extended Data Fig. 9. Nutrient starvation induces SCARF via eIF5B degradation.**

**(a)** Histogram showing the fold change of ribosome density after amino acid starvation. The mRNAs with highest fold change values (top 25%) were defined as Inc, and highlighted by red. The mRNAs with lowest fold change values (bottom 25%) were defined as Dec, and highlighted by blue. The right panels show IFR along CDS of the two groups of mRNAs. IFR values are calculated within a non-overlapping sliding window (45 nt), which was subsequently normalized by CDS IFR.

**(b)** Bar graphs show the HiBit-based SCARF reporter assays in cells before and after amino acid starvation. Both the Fluc (left panel) and HiBit (right panel) signals were measured from cells transfected with SCARF reporters bearing a uORF start codon with or without the Kozak sequence context. Error bars, mean ± s.e.m.; two-tailed *t*-test, $n = 3$, *** $P < 0.001$.

**(c)** The top panel shows the half-life of eIF5B in HEK293 cells before and after amino acid starvation. Cells were treated with cycloheximide (100 μg/mL) for various times followed by immunoblotting of whole cell lysates. The bottom panel shows the half-life of eIF5B in starved HEK293 cells in the presence of 5 μM MG132.

**(d)** Representative flow cytometry scatter plots of HEK293-$K^b$ cells transfected with SCARF reporters before and after amino acid starvation, in the absence or presence of 5 μM MG132. Bar plots show the relative 25D1 mean fluorescence intensity (MFI) of SCARF reporters. The highlighted numbers refer to the reading frame of the encoded SIINFEKL relative to the AUG codon. Error bars, mean ± s.e.m.; two-tailed *t*-test, $n = 3$, *** $P < 0.001$.

**(e)** Western blots of exogenous eIF5B in transfected HEK293 cells.

**(f)** Representative flow cytometry scatter plots of HEK293-$K^b$ cells transfected with SCARF reporters before and after amino acid starvation, in the absence or presence of exogenous eIF5B. Bar plots show the relative 25D1 mean fluorescence intensity (MFI) of SCARF reporters. The highlighted numbers refer to the reading frame of the encoded SIINFEKL relative to the AUG codon. Error bars, mean ± s.e.m.; two-tailed *t*-test, $n = 3$, *** $P < 0.001$.

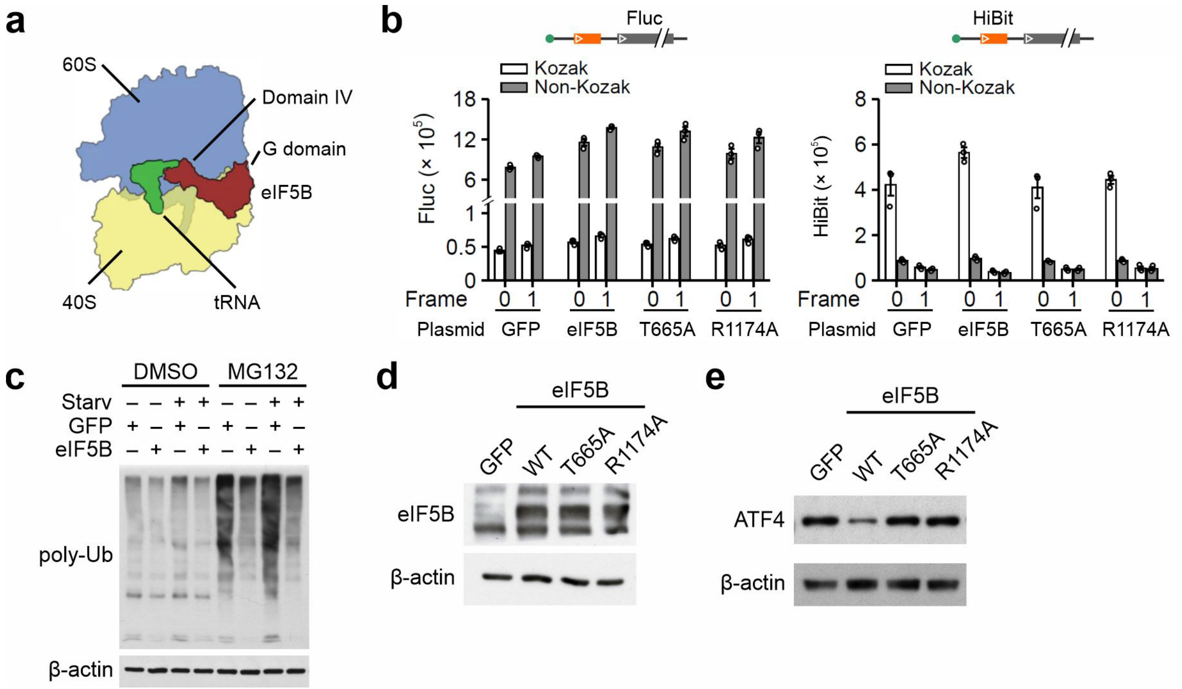


**Extended Data Fig. 10. Rescuing effect of eIF5B in nutrient starvation induced SCARF.**

**(a)** A schematic image of 80S-eIF5B complex adopted from Wang *et al*, Nat Commun 2020.

**(b)** Bar graphs show the HiBit-based SCARF reporter assays in cells transfected with wild type eIF5B or mutants. Both the Fluc (left panel) and HiBit (middle panel) signals were measured from cells transfected with SCARF reporters bearing a uORF start codon with or without the Kozak sequence context.

**(c)** Western blots of polyubiquitinated species in HEK293 cells with or without eIF5B overexpression before and after amino acid starvation.

**(d)** Western blots of exogenous eIF5B in transfected HEK293 cells.

**(e)** Western blots of ATF4 in starved HEK293 cells transfected with eIF5B wild type or mutants.
